# Rats in the city: implications for zoonotic disease risk in an urbanizing world

**DOI:** 10.1101/2021.03.18.436089

**Authors:** Kim R. Blasdell, Serge Morand, Susan G.W. Laurance, Stephen L Doggett, Amy Hahs, David Perera, Cadhla Firth

## Abstract

Urbanization is rapidly transforming much of Southeast Asia, altering the structure and function of the landscape, as well as the frequency and intensity of the interactions between people, animals, and the environment. In this study, we began to explore the impact of urbanization on zoonotic disease risk by simultaneously characterizing changes in the abundance and diversity of reservoir hosts (rodents), ectoparasite vectors (ticks), and microbial pathogens across a gradient of urbanization in Malaysian Borneo. We found that although rodent species diversity decreased with increasing urbanization, two species appeared to thrive in anthropogenic environments: the invasive urban exploiter, *Rattus rattus* and the native urban adapter, *Sundamys muelleri*. *R. rattus* was strongly associated with the presence of built infrastructure across the gradient and dominated the urban rodent community where it was associated with high microbial diversity and multi-host zoonoses capable of environmental transmission, including *Leptospira* spp., and *Toxoplasma gondii*. In contrast, *S. muelleri* was restricted to sites with a significant vegetative component where it was found at high densities in the urban location. This species was strongly associated with the presence of ticks, including the medically important genera *Ambylomma*, *Haemaphysalis*, and *Ixodes*. Overall, our results demonstrate that the response to urbanization varies by species at all levels: host, ectoparasite, and microbe. This may lead to increased zoonotic disease risk in a subset of environments across urban and urbanizing landscapes that can be reduced through improved pest management and public health messaging.

## Introduction

Urbanization is a widespread and significant process of global change that modifies the landscape rapidly, extensively, and often permanently. These environmental changes are accompanied by a marked reduction in biodiversity in cities that is driven by varied responses from plant and animal species (Grimm et al. 2008). Many species are highly sensitive to the effects of urbanization, including the loss or fragmentation of suitable habitat, and may disappear from the urban environment completely. However, the process of urbanization also creates new environmental conditions and an abundance of novel resources. These enable some wildlife (termed urban exploiters), to be highly successful in cities, where they are often found at unusually high densities (Shochat et al. 2006, McFarlane et al. 2012). Within mammals, characteristics of urban exploiters include large litter sizes, as well as traits associated with behavioural plasticity and generalist diets (Santini et al. 2019). These characteristics allow urban mammals to exploit the new habitats and resources available in city environments, while adapting to and avoiding the many risks of urban living.

Several rodent species are amongst the most successful urban dwellers. These include the ubiquitous commensals, *Mus musculus* (the house mouse)*, Rattus norvegicus* (the Norway rat), and the *Rattus rattus* species complex (the black rat and its relatives), which are thought to have coexisted with people for thousands of years. Recently it has been argued that these urban exploiters are now so well-adapted to cities that they should be considered native wherever they are found in the urban environment, irrespective of the location of the city (Banks and Smith, 2015). As urbanization has intensified globally, an increasing number of native rodents also appear to be effectively adapting to local city environments (termed ‘urban adapters’), where they are frequently found within both planned and remnant green spaces (Baker et al. 2003, Banks and Smith 2015, Castillo et al. 2003, Parsons et al. 2018).

Unfortunately, rodent presence in cities can be both an economic burden (rodent infestation costs an estimated ∼$19 billion in damage to infrastructure and food supplies in the US each year), and a threat to human health (Pimentel et al. 2000, Himsworth et al. 2013). Urban rodent infestations have been associated with an increased risk of asthma and allergies, as well as with the spread of a range of zoonotic and food-borne illnesses, including leptospirosis, hantavirus, Lassa fever virus, *Salmonella enterica*, and vector-borne bacterial infections, including scrub typhus (Berg et al. 2007, Blanco Crivelli et al, 2012, Blasdell et al. 2019b, Bonell et al. 2017, Chew et al. 2003, Himsworth et al. 2013, Firth et al. 2014). However, little is known about the drivers of rodent-borne zoonotic disease transmission in cities, or how we might reduce the impact of urban rodents on human health through changes to infrastructure, resource distribution, or human behavior.

Tropical cities often bear the brunt of rodent-associated problems, with informal infrastructure and limited sanitation combining to create favorable environments for rodents, while warmer temperatures and increased humidity act to improve the environmental survival and transmissibility of some of the pathogens they carry (Bonell et al. 2017, Costa et al. 2015, Meerburg et al. 2009). In addition, many arthropod vectors of rodent-borne diseases thrive under tropical conditions, including ticks, mites, and mosquitoes, which may further increase human disease risk. Critically, ∼90% of urban population growth in the next 30 years is also expected to occur in lower- and middle-income countries in tropical and sub-tropical Asia and Africa (United Nations, 2014). This rapid growth has the potential to exacerbate many of the public and environmental health challenges that exist in these regions, including those posed by urban rodents. Unfortunately, research into the ecology of urban rodents and associated zoonoses has been biased towards temperate cities, with preliminary investigations into rodent-borne pathogens in the tropics available for only a handful of cities, including Bangkok, Kuala Lumpur, Morogogo, and Salvador (Tantivanich et al. 1992, Siritantikorn et al. 2003, Mgode et al. 2005, Günther et al. 2008, Paramasvaran et al. 2009, Benacer et al. 2013, Costa et al. 2014, Minter et al. 2017, Walker et al. 2017, Premaalatha et al. 2018, Tan et al. 2019).

Despite the increasing relevance of urban ecosystems to human health in the tropics, little is known about how rodents use these urban and urbanizing landscapes, the features of the environment that promote their persistence, or how the process of urbanization is likely to affect human disease risk. In this study, we explored how the abundance and diversity of rodents and their ectoparasites varied across a gradient of urbanization surrounding a tropical city in Malaysian Borneo (Kuching, Sarawak), and examined the environmental features associated with their presence in urban and urbanizing environments. To begin to understand how urbanization might impact zoonotic disease risk in tropical cities, we further examined the prevalence of select groups of rodent-borne microbes across the urban-rural gradient, and discuss how urbanization may affect human disease risk.

## Methods

### Ethics statement

This study was approved by the CSIRO Australian Animal Health Laboratory’s Animal Ethics Committee (#1750) and the Sarawak Forests Department (Permit: NCCD.907.4.4 (JLD.12)-131).

### Rodent and environmental data collection

Rodents were collected at three locations representing a gradient of urbanization, including: (1) “Mount Singai” – a rural location with low levels of urbanization, dispersed villages and semi-disturbed vegetation; (2) “Batu Kawa district” – a rapidly developing region on the outskirts of Kuching with moderate levels of urbanization, disturbed vegetation, and both new and old human settlements; and (3) “Central Kuching” – which is comprised of urban grey/green matrix at the center of Sarawak’s largest city, and is located at 1.54°N, 110.35°E (Figure 1). A detailed description of each location is provided in Blasdell et al. (2019b). For clarity, locations will be referred to as ‘rural, ‘developing’, and ‘urban’ throughout the manuscript.

**Figure 1.**
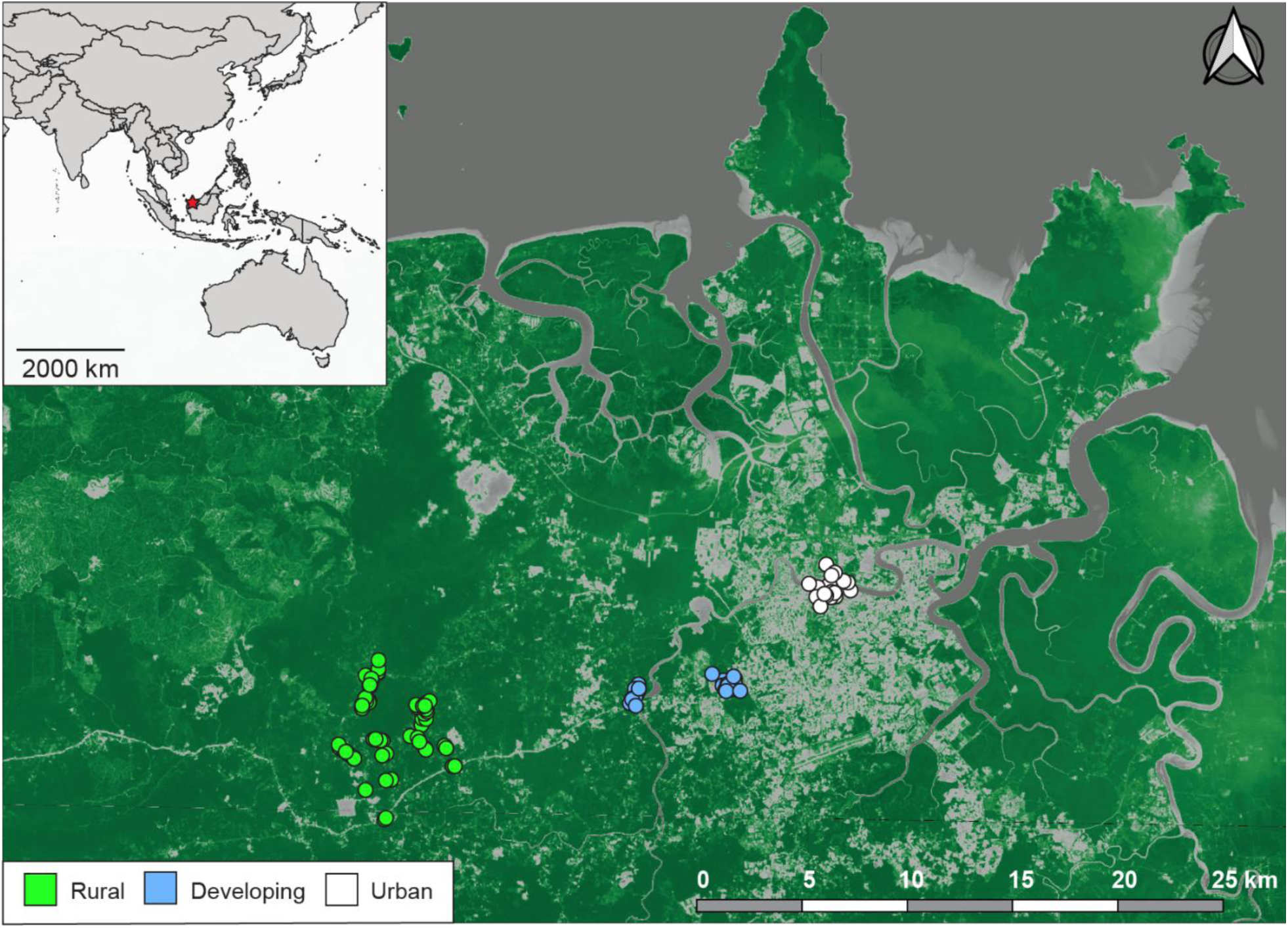
Sites of rodent collection across the urban-rural gradient. From left-to-right: rural (green circles; Mount Singai), developing (blue circles; Batu Kawa district), and urban (white circles; Central Kuching) locations are shown. The inset in the upper-left corner indicates the location of the study – Sarawak, Malaysia with the city of Kuching marked by a red star.

Rodent trapping within each location was conducted at multiple sites, which were defined as a circle of 110m radius centered at the point where GPS coordinates were taken (Figure 1). The 110m radius was chosen to correspond with the approximate home range of *R. rattus* estimated under similar environmental conditions, as this was the only study species for which suitable home range data was available (Harper et al. 2015). Sites were categorized based on the dominant land-use type observed within the complete circle (i.e., commercial, forest, residential/village, scrub and mixed commercial/residential), and if a site was on (or next to) an ecotone, this was also noted. A number of site-specific environmental variables were measured or estimated by ground truthing at the time of rodent collection, including the presence of: (i) waterbodies (e.g., river, stream, swamp, pond); (ii) sewers; (iii) livestock (e.g., chickens, pigs, other); (iv) food plants (e.g., banana, corn, sugar); (v) rubbish; (vi) roads; (vii) buildings and building condition; (viii) commercial food storage/service/preparation areas, and (ix) green space (Supplemental Table 1).

Rodent trapping was conducted across five time intervals that bracketed the wet season: (1) September/October 2015, (2) March/April 2016, (3) September/October 2016, (4) March/April 2017, (5) September/October 2017. Rather than equalizing trapping effort at each location, we attempted to collect a minimum of 50 rodents per location per time, which resulted in sampling from a total of 27 urban, 23 developing, and 65 rural sites across the study period (N=115 total) (Figure 1). Trapping was conducted at each site for between one and seven nights using locally-made wire mesh traps (∼30cm x 14cm x 14cm), baited daily with meat and fruit, and placed at intervals >1.5m. Traps were opened between 4 pm - 6pm daily and collected/closed between 6am - 8am the following morning. At the time of collection, rodents were morphologically assigned a species identification and either immediately released (N=47), or euthanized by over-anaesthetization in isoflurane followed by bilacteral thoracotomy (N=815). For those animals that were euthanized, carcasses were fumigated using ethyl acetate for ∼5 min to kill ectoparasites and combed with a fine-toothed flea comb. Ectoparasites were sorted visually by category (i.e., mites, fleas, lice, ticks), and placed either in ethyl alcohol for further identification, or frozen directly on dry ice. Ticks were identified using the taxonomic keys and texts of Durden et al. (2008), Keirans et al. (1970), Kohls (1957), Lah et al. (2016) and Madinah et al. (2013). However, we were only able to identify ticks to the genus level due to the lack of larval and nymphal keys, as these life stages comprised the vast majority of our collection. Lice (Phthiraptera) and fleas (Siphonaptera) were identified only to order, and mites were only identified to the subclass Acari. The weight, sex, and reproductive status of each animal was recorded, and tissue samples (blood, ear, lung, kidney, liver, spleen) were aseptically collected and frozen directly on dry ice prior to storage at −80°C. The complete data set associated with this publication is available on DataDryad.org

### Landcover estimates

Site-specific landcover estimates were calculated in QGIS v3.2.3 by first using the Semi-Automatic Classification Plugin v6.2.9 to transform LANDSAT 8 data (https://earthexplorer.usgs.gov/) into reflectance values (Congedo 2016, QGIS Development Team (2019). These values were used to calculate the normalized difference vegetation index (NDVI) for each site using the QGIS Raster Calculator tool and the equation: NDVI = (Band 5 – Band 4) / (Band 5 + Band 4), where Band 4 = red and Band 5 = near-infrared. Pixels with NDVI values of −1.000 – 0.150 were considered water, 0.151 - 0.700 were considered urban (grey), and 0.701 – 1.000 were considered green. For downstream analyses, we calculated: (i) NDVI_mean as an estimate of the average proportion of green landcover at each site; (ii) NDVI_stdev as an estimate of landcover variability across a site; (iii) distance_to_urban, the distance from the center of each site to the nearest ‘urban’ pixel (an indicator of access to resources associated with urban landcover); and (iv) distance_to_water, the distance from the center of each site to the nearest ‘water’ pixel. Finally, we explored using datasets from the Global Road Inventory Project (GRIP); https://datacatalog.worldbank.org/dataset/grip-global-roads-inventory-project-2018), the Global Urban Landcover dataset (ftp://115.239.252.28/GlobalUrbanLand/), and the WorldPop demographic dataset (https://www.worldpop.org/) to calculate additional estimates of urbanization; however, none of these were of appropriate resolution for the study area.

### Molecular analyses

DNA and RNA were extracted from ∼30mg of homogenized tissue using the AllPrep DNA/RNA Mini kit (Qiagen, Valencia, CA) as per the manufacturer’s instructions. Nucleic acid quality and concentration for each sample were assessed using the NanoDrop-1000 spectrophotometer (Thermo Scientific, Somerset, NJ), and extractions were repeated for samples with a 260/280 < 1.8 or that yielded < 50ng/ul of nucleic acid. cDNA synthesis was performed on each RNA extraction using Superscript III (Invitrogen, Carlsbad, CA), 50ng of random primers, and 50-200 ng of template RNA, following the manufacturer’s protocol. The tentative species assignment given to each animal at the time of collection was confirmed by sequencing the product of a polymerase chain reaction (PCR) assay using primers BatL5310 and R6036R to amplify ∼750bp of the cytochrome oxidase I (COI) gene (Robins et al. 2007). This was followed by BLAST® analysis (http://blast.ncbi.nlm.nih.gov/Blast.cgi) to assess sequence similarity and assign a species identification.

A subset of animals (N=316) was selected for additional molecular screening to identify common rodent-borne microbes, many of which are zoonotic pathogens. These were comprised of individuals belonging to the species *Maxomys whiteheadi* (N=12), *M. ochraceiventer* (N=2), *Niviventer cremoriventer* (N=14), *Niviventer* sp. (N=1), *Rattus rattus* lineage R3 (N=165), *R. rattus* lineage *tanezumi* (N=11), *R. tiomanicus* (N=8), *R. exulans* (N=3) and *Sundamys muelleri* (N=100), and were all collected between September 2015 and April 2016. Previously published PCR assays were used to screen appropriate tissue and nucleic acid types for selected viruses: alphaviruses, flaviviruses, enteroviruses, mammarenaviruses, coronaviruses, orthohantaviruses, orthohepeviruses, paramyxoviruses; bacteria: *Bartonella* spp., *Leptospira* spp., *Rickettsia* spp. *Yersinia pestis*; and the intracellular parasite, *Toxoplasma gondii* (see Supplemental Methods for details of tissues and assays). PCR products with appropriately-sized amplicons in any assay were subjected to Sanger sequencing, followed by BLAST® analysis to assess sequence similarity. Multiple sequence alignments and phylogenetic trees were constructed when further sequence analysis was required, and were performed using MUSCLE v3.8 and PhyML v3.1, respectively (Edgar 2004, Guindon et al. 2010).

### Statistical analyses

Rodent species richness was estimated using the first-order jackknife (Jack1) and bootstrap method estimators in the package ‘vegan’, implemented in R (Oksanen et al. 2019, R Core Team 2018). Rarefaction curves were used to estimate the number of rodent species present (as a function of the number of sites), for each location. Estimates of Beta diversity (ß) were also calculated in ‘vegan’ using the Bray-Curtis index. Values of ß range from 0 (the community composition at two locations are completely dissimilar), to 1 (community composition at two locations are identical).

Two Linear Discriminant Analyses (LDA) were used to assess the fit of our *a priori* assignment of each site into a ‘rural’, ‘developing’, or ‘urban’ classification using a subset of environmental variables. The first LDA (LDA_strict) contained only the landscape variables: NDVI_mean, NDVI_stdev, distance_to_urban, and distance_to_water. Subsequently (LDA_relaxed), we added site-specific variables related to the presence/absence of livestock, sewers and buildings. LDAs were performed using the library MASS, implemented in R (Venables and Ripley 2002).

To identify links between rodent presence and the environmental variables measured across the urban-rural gradient, non-metric multidimensional scaling (NMDS) was first used to ordinate all 115 sites in environmental space using PC-ORD 6.08 (McCune and Mefford 2011). For this analysis, a Sørensen dissimilarity matrix was calculated based only on those environmental variables for which non-zero values were recorded for at least 50% of sites. These included: waterbody (flowing, standing, none); sewers (presence/absence); chickens (presence/absence); food plants (presence/absence); rubbish (new/old/none); roads (presence/absence); dominant green space type (forest, scrub, mixed, garden, none), and the fraction of each site that was ‘green space’ or ‘buildings’, for a total of nine variables. A Monte Carlo randomization test was applied to determine the stress of the final solution. The resulting ordination axes were correlated with each environmental variable using the Pearson correlation coefficient with a Bonferroni correction for multiple tests to establish a conservative threshold for significance (P < 0.005, n = 115 sites). Finally, these ordination axes were used as predictor variables in independent binomial generalized linear models (GLMs) to examine their influence on rodent presence. GLMs were constructed individually for the two most frequently collected rodent species (*R. rattus* and *S. muelleri*), using the R packages ‘car’ and ‘ResourceSelection’ (Fox & Weisberg 2019, Lele et al. 2019). Model fit was assessed using the Hosmer-Lemeshow goodness of fit test. As a comparison, GLMs were also constructed for both *R. rattus* and *S. muelleri* as above, with ‘location’ (rural, developing, urban) as a predictor.

Descriptive statistics (Fisher’s exact test, Chi-square test) were used to investigate the relationships between ectoparasite presence/abundance (ticks, lice, mites) and location (rural, developing, urban), as well as associations with rodent species. Exact goodness of fit tests were used to assess differences in trapping success by sex, and one-way ANOVAs followed by paired t-tests (with Bonferroni correction for multiple comparisons) were used to examine changes in adult body mass index (calculated as the ratio: body weight (g) / snout-vent length (cm)) between locations. No pregnant individuals were included. We focused solely on the two most abundant rodent species for these analyses (*R. rattus* and *S. muelleri*) due to small sample sizes of the remaining species.

Bipartite networks were constructed to examine the interactions between: (i) ticks and rodents, and (ii) microbes and rodents, at each location across the urban-rural gradient using the R packages ‘bipartite’ v2.03 and ‘igraph’ (Dormann et al. 2009, Csardi and Nepusz 2006). The tick community was described by both genus (*Amblyomma*, *Haemaphysalis*, or *Ixodes*) and life stage (larva, nymph, adult), while microbes and the rodent community were described at the species level. Interactions between rodents and either ticks or microbes were investigated using three qualitative metrics. First, the connectance of each network was calculated as the ratio of observed to possible links between rodents and parasites, which corresponds to the number of species combinations in the system (Dunne et al. 2002). Next, we calculated the modularity of each network to identify subsets (modules) of the network that are more densely connected than other subsets. Higher values of modularity indicate that subsets of rodents and parasites interact more strongly amongst themselves (i.e, within a module), than with other subsets (Thébaud 2013). Finally, we calculated the nestedness temperature of each network, which provides an estimate of the orderliness of the network and reflects both species composition and incidence (Ulrich et al. 2009). Nestedness temperature values range from 0 to 100, with higher values observed when more abundant parasites interact more frequently with a wider range of hosts. Finally, we transformed each bipartite network into a unipartite network by connecting individual rodents based on shared ticks/microbes using the ‘igraph’ package in R (Csardi and Nepusz 2006). Nonparametric resampling was used to estimate the edge density with confidence intervals of each unipartite network using the R package ‘snowboot’ (Ramirez-Ramirez et al. 2019). Eigenvector centrality was calculated from each projected unipartite network to identify individuals, collection sites, or rodent species occupying central (i.e., well-connected) positions in each network, and which may be of importance for the transmission of ticks or microbes at each location.

## Results

Our field classification of the 115 study sites as rural, developing, and urban concurred with a GIS-based classification for 85.2% (98/115) of sites when only four environmental variables were included (LDA_strict, Supplemental Figure 1). When additional variables that described the presence of livestock, sewers and buildings were added (LDA_relaxed, Supplemental Figure 1), the proportion of correctly classified sites improved to 89.6% (103/115). Rural sites were most likely to be correctly classified (60/65 sites correctly classified by LDA_strict, 63/65 sites by LDA_relaxed), with all incorrectly predicted sites classified as developing. In each of these cases, misclassified sites contained significant anthropogenic influence and incorporated the edge of a village or multi-family dwellings. In contrast, urban sites were most likely to be incorrectly classified (20/27 sites correctly classified by by LDA_strict, 21/27 by LDA_relaxed), and again, each mis-assigned site was classified as developing. Similarly, 18/23 (LDA_strict) and 19/23 (LDA_relaxed) developing sites were correctly classified; however, incorrectly predicted sites were assigned to both urban and rural locations.

### Rodents and the urban-rural gradient

Across all sites, 863 rodents were captured over 7665 trap-nights. We collected 246 rodents from the rural location, 318 from the developing location, and 299 from the urban location (Supplemental Table 2). These were comprised of Individuals from ten species, with the majority (92%) assigned to the species *R. rattus* lineage R3 (N=375), *S. muelleri* (N=331), *R. tiomanicus* (N=54), and *R. rattus* lineage *tanezumi* (N=41). These four species were also the only species that were collected over the complete urban-rural gradient. There was a clear decrease in rodent species richness across the gradient; ten species were sampled in the rural location (Jack1 estimator = 16.7 ± 8.0; bootstrap estimator = 20.0 ± 12.9), seven in the developing location (Jack1 estimator = 11.7 ± 5.7; bootstrap estimator = 9.1 ± 3.2), and four in the urban location (Jack1 estimator = 6.7 ± 3.1; bootstrap estimator = 5.2 ± 1.8). This trend was also obvious when species diversity was normalized by sample size (Figure 2). However, rodent community composition was highly similar between the rural and developing locations (ß = 0.824), with decreasing similarity between developing and urban locations (ß = 0.730), and urban and rural locations (ß = 0.571). Across all rodent species, slightly more males (51.9%) than females were trapped; however, these differences were not significant for any of the four most abundant species collected (Supplemental Table 1, P > 0.05). For all species, the vast majority of rodents captured were adults, with the highest proportion of juvenile animals belonging to the species *R. rattus* lineage *tanezumi* (14.6%, Supplemental Table 1).

**Figure 2.**
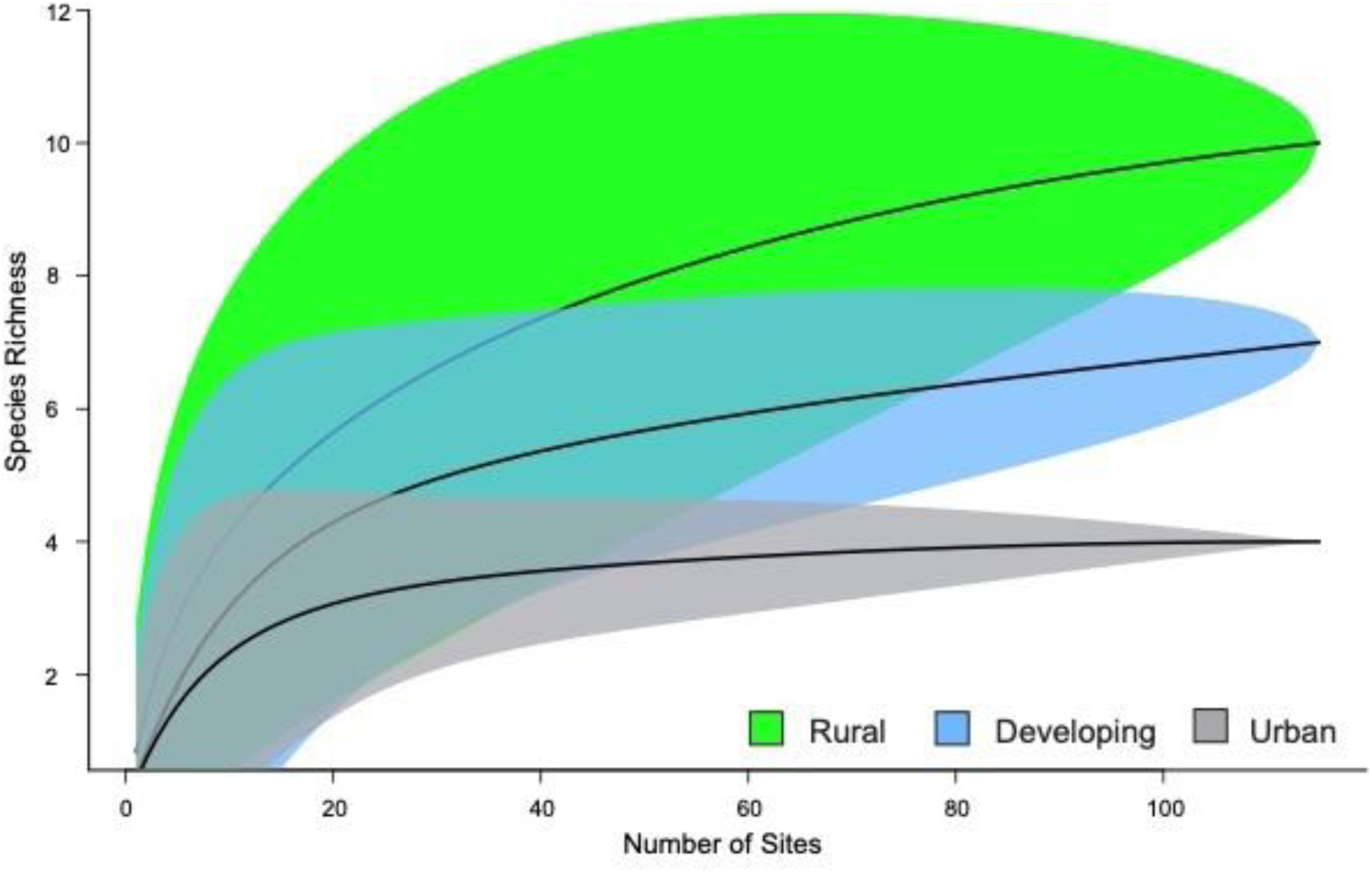
Rodent species richness rarefaction curves across the rural-urban gradient. Expected number of rodent species at each location is shown as a function of the number of sites sampled, with 95% confidence intervals.

### Response to urbanization: R. rattus and S. muelleri

To further explore the impact of urbanization on the distribution of rodents, we focused more detailed analyses solely on the two most abundant rodent species across the urban-rural gradient: *R. rattus* and *S. muelleri*. Two distinct mitochondrial lineages of the *R. rattus* species complex were identified via the COI barcoding PCR: *R. rattus* lineage R3 and *R. rattus* lineage *tanezumi*. As multiple lines of evidence now suggest that these two lineages should both be considered *R. rattus,* we considered them collectively for all further analyses, unless otherwise specified (Musser & Carleton 2005; Pages et al. 2010, 2013, Aplin 2011, Wells et al. 2014, Lau et al. 2020).

Across the urban-rural gradient, 73% of all sites contained *R. rattus*, while 44% of sites contained *S. muelleri*. The proportion of sites with *R. rattus* increased with increasing anthropogenic influence across the gradient, whereas *S. muelleri* was most common at sites in the developing location (Figure 3). The proportion of sites at which both species were collected also increased across the urban-rural gradient, with significantly more shared sites in the urban and developing locations (47.4% and 41.7% of sites shared, respectively), than in the rural location (16.9%; P = 0.01). Notably, we found that location alone was not a strong predictor of either *R. rattus* or *S. muelleri* presence at a site (Table 1). As a result, we ordinated our study sites using NMDS and nine environmental variables, which resolved highly orthogonal principal axes in three dimensions. These three axes explained 86.3% of the variation in our data, and the stress of the final solution was less than 99.3% (P<0.004). Strikingly, sites were only slightly distributed across these axes according to location (rural, developing, urban), which suggests that these categories alone are not sufficiently able to capture the fine-scale environmental gradients present across the landscape (Figure 3). GLMs were constructed to explore the association between *R. rattus* and *S. muelleri* and the three ordination axes, which revealed a highly significant relationship between the presence of *R. rattus* and Axes 1 and 2, and a significant relationship between the presence of *S. muelleri* and Axes 2 and 3 (Table 1, Figure 3).

**Figure 3.**
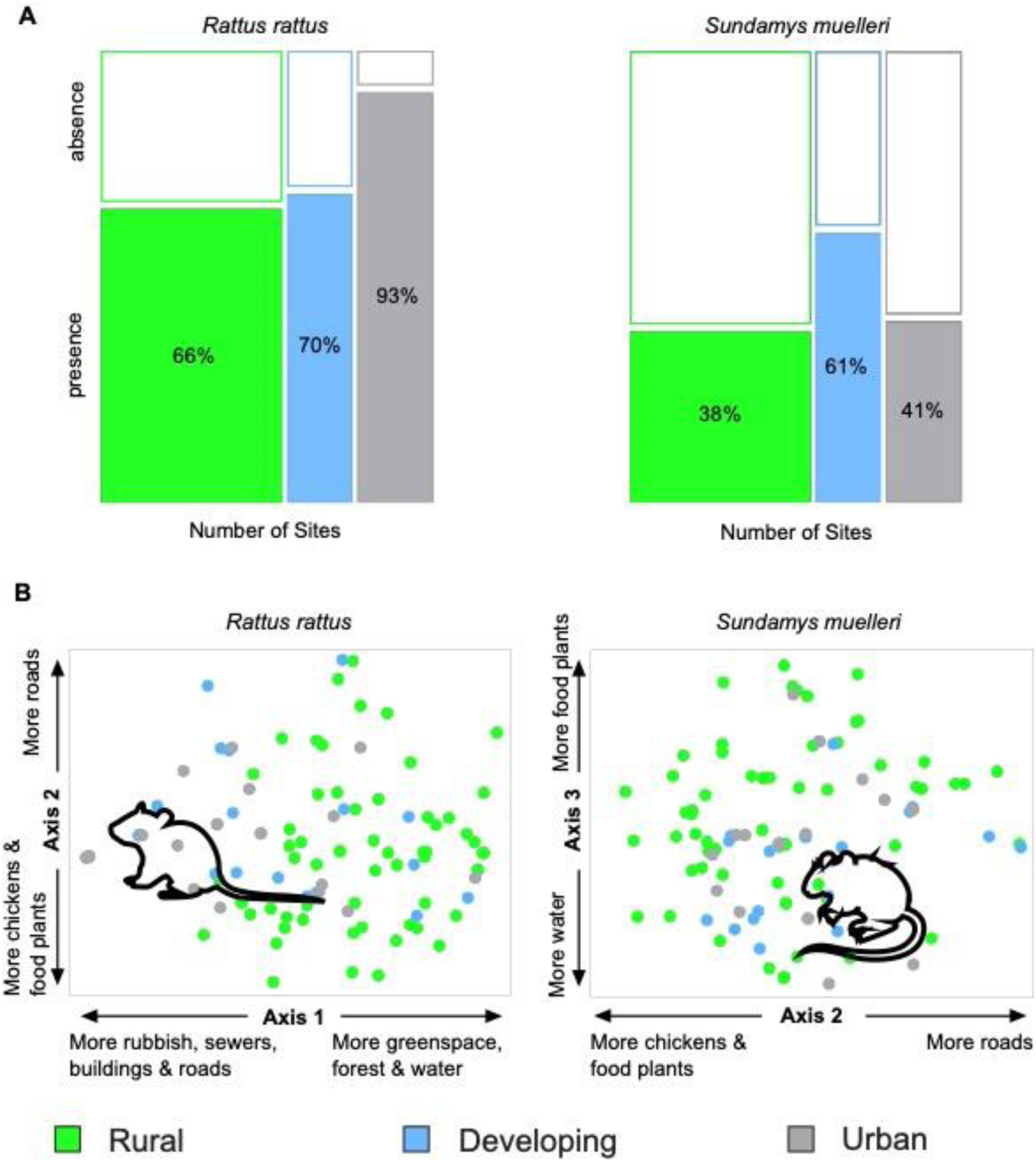
Distribution of *R. rattus* and S. *muelleri* across the urban-rural gradient and the environmental factors that best predict their presence. **Panel A**: the proportion of sites positive for each species is shown on the vertical axis with the number of sites sampled represented on the horizontal axis. **Panel B**: The distribution of rural, developing and urban sites is shown according to the three principal axes ordinated by NMDS. The environmental features most likely to be associated with each species is indicated by the position of the rodent symbol.

**Table 1.**
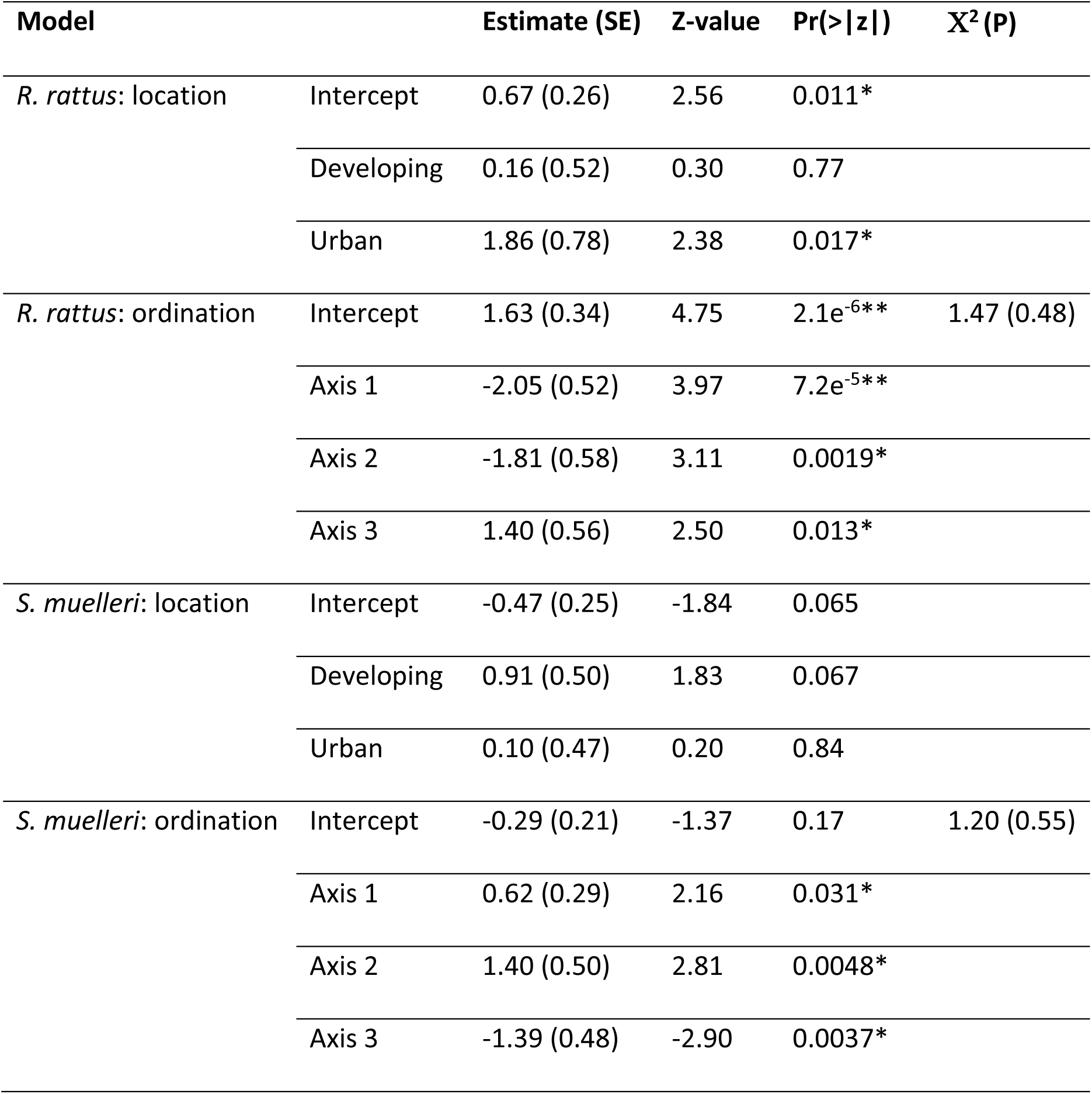
Results of the binomial GLMs that describe the presence of R. rattus and S. muelleri across the urban-rural gradient. Values are shown for (i) estimated regression coefficients (Estimate) with standard error (SE), (ii) Z-values, and (iii) P-values (Pr(>|z|). Significance at α = 0.05 is indicated by *; significance at α = 0.001 is indicated by **. The value for the Hosmer-Lemeshow Chi-square (X^2^) and associated P=value (P) is also given for each best model.

| Model | | Estimate (SE) | Z-value | $\Pr(>\vert z\vert )$ | $X^2$ (P) |
| --- | --- | --- | --- | --- | --- |
| *R. rattus*: location | Intercept | 0.67 (0.26) | 2.56 | 0.011* |  |
|  | Developing | 0.16 (0.52) | 0.30 | 0.77 |  |
|  | Urban | 1.86 (0.78) | 2.38 | 0.017* |  |
| *R. rattus*: ordination | Intercept | 1.63 (0.34) | 4.75 | $2.1e^{-6**}$ | 1.47 (0.48) |
| | Axis 1 | -2.05 (0.52) | 3.97 | $7.2e^{-5**}$ | |
|  | Axis 2 | -1.81 (0.58) | 3.11 | 0.0019* |  |
|  | Axis 3 | 1.40 (0.56) | 2.50 | 0.013* |  |
| *S. muelleri*: location | Intercept | -0.47 (0.25) | -1.84 | 0.065 |  |
|  | Developing | 0.91 (0.50) | 1.83 | 0.067 |  |
|  | Urban | 0.10 (0.47) | 0.20 | 0.84 |  |
| *S. muelleri*: ordination | Intercept | -0.29 (0.21) | -1.37 | 0.17 | 1.20 (0.55) |
|  | Axis 1 | 0.62 (0.29) | 2.16 | 0.031* |  |
|  | Axis 2 | 1.40 (0.50) | 2.81 | 0.0048* |  |
|  | Axis 3 | -1.39 (0.48) | -2.90 | 0.0037* |  |

Axis 1 explained the majority of the variation in our environmental data (R^2^=0.587), and distributed the sites based on a gradient of human infrastructure, while Axis 2 (R^2^=0.157) reflected a gradient of small-scale domestic agriculture (Table 2, Figure 3). Axis 3 explained the least amount of data variation (R^2^=0.119), and placed the sites on a gradient primarily driven by the presence of natural waterbodies. Taken together, these results suggest that *R. rattus* was significantly more likely to be found at sites with built infrastructure, including buildings, roads, rubbish, and sewers, while *S. muelleri* was more likely to be found at green sites with flowing water and without food plants or chickens (Figure 3). However, due to the smaller number of *S. muelleri* sampled (and smaller number of sites with *S. muelleri* present), we have less statistical confidence in our ability to predict the presence of this species based on the three ordination axes (Table 1).

**Table 2.** Pearson correlation coefficients of environmental variables with ordination axes. Significance is indicated as follows: * P<0.005; ** P<0.0001; *** P<0.00001.

| Variable | Ordination Axis 1 | Ordination Axis 2 | Ordination Axis 3 |
| --- | --- | --- | --- |
| Fraction Buildings | -0.715*** | -0.240 | 0.115 |
| Chickens | 0.362* | -0.299* | 0.274 |
| Food-plants | 0.340* | -0.390** | 0.441** |
| Greenspace Type | 0.549*** | 0.093 | 0.088 |
| Fraction Greenspace | 0.764*** | -0.100 | 0.227 |
| Roads | -0.511*** | 0.379** | -0.020 |
| Rubbish | -0.807*** | -0.309* | 0.018 |
| Sewers | -0.738*** | -0.152 | 0.294* |
| Waterbody | 0.553*** | -0.522*** | -0.647*** |

Finally, we examined the body condition (i.e., body mass index) of adult *R. rattus* and *S. muelleri* at each location as a proxy for the health of individuals at the time of sampling. Body condition was significantly different between male and female individuals for both species across the urban-rural gradient; as a result, each sex was analyzed separately (P < 0.05) (Figure 4). Overall, the body mass index of individuals from both species varied significantly by location (P < 0.002) for all sex and species combinations. Pairwise comparisons revealed that for both male and female *R. rattus* and *S. muelleri*, individuals from the urban location were significantly heavier/healthier than those from the rural location (P < 0.0167). Similarly, male *S. muelleri* and both male and female *R. rattus* from developing areas were significantly heavier than individuals from the rural location, while female *S. muelleri* from the urban location were significantly heavier than those from the developing location (P < 0.0167). There was no statistical difference in body condition between male/female *R. rattus* or male *S. muelleri* from the developing and urban locations, nor between female *S. muelleri* from the rural and developing locations. There was also no significant difference in the number of pregnant and non-pregnant females collected at each location.

**Figure 4.**
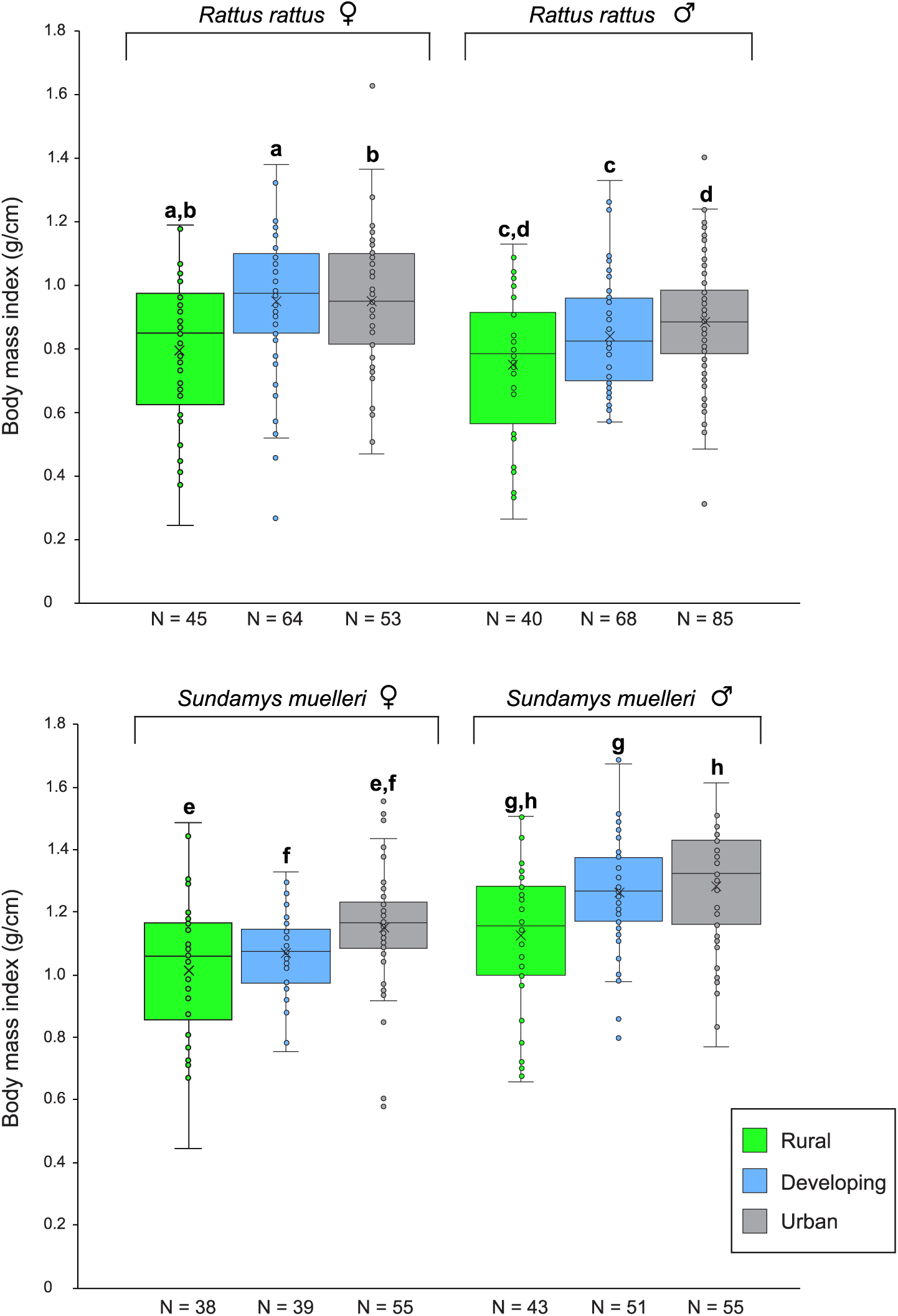
Adult body condition of *R. rattus* and *S. muelleri* across the urban-rural gradient. Body condition was estimated by calculating the body mass index of male and female *R. rattus* and *S. muelleri* as follows: (weight (g) / snout-vent length (cm)). For each comparison, significant differences in body condition between locations are indicate by lower case letters (corrected α = 0.0167).

### Ectoparasites and the urban-rural gradient

Of the 815 animals from which ectoparasites were collected, only 105 rodents (12.8%) were free of any ectoparasite. Mites (Mesostigmata) were most commonly identified, followed by ticks (Ixodida), lice (Phthiraptera), and fleas (Siphonaptera) (Table 3). Co-infections were routinely identified, with 188 (23.1%) animals infested by at least two groups of ectoparasite. Of the two most abundant rodent species (*S. muelleri* and *R. rattus*), mites were significantly more likely to be found on *S. muelleri* (Fisher exact test, P < 0.01), where they were also more abundant on each individual (personal observation). Three genera of hard ticks (Family *Ixodidae*) were identified: *Amblyomma* spp., *Haemaphysalis* spp., and *Ixodes* spp. All three life stages (i.e., larvae, nymphs, adults) of *Amblyomma* ticks were present only on *R. rattus*, whereas all three life stages of *Ixodes* ticks were only collected from *S. muelleri* (Supplemental Table 3). Twenty-six individuals from five species of rodents were parasitized by adult *Ixodes* ticks, none of which were collected from the urban location. No adult *Haemaphysalis* ticks were present on any rodent sampled, and only a single adult *Amblyomma* was encountered. All three tick genera were sampled from the same individual on three occasions, and 39 individuals were infested with more than five ticks, although most ticks were encountered as singletons. All tick genera were more strongly associated with *S. muelleri* than *R. rattus*, with significantly more *S. muelleri* individuals infested (Fisher exact test, P < 0.01). This pattern was also observed when each tick life stage was examined separately: *S. muelleri* was significantly more likely to be infested with *Haemaphysalis* and *Amblyomma* larvae, all three genera of nymphs (Fisher exact test, P < 0.01), and *Ixodes* adults. Although *S. muelleri* was also more commonly parasitized by *Ixodes* larvae than *R. rattus*, this result was not significant (P < 0.10). Finally, individual *S. muelleri* were the most heavily parasitized species in this study; of rodents with at least one tick present (N = 144), 70% belonged to *S. muelleri,* as did 14 of the 15 individuals with more than ten ticks.

**Table 3.** Ectoparasites collected from each rodent species. For each species, the number and proportion (%) of rodents infested are shown. Zeros indicate no ectoparasite was found, “-“ indicates no animals were sampled. R = Rural location, D = Developing location, U = Urban location, T = Total.

| Species<br>(N sampled) | Fleas |  |  |  | Lice |  |  |  | Mites |  |  |  | Ticks |  |  |  |
| --- | --- | --- | --- | --- | --- | --- | --- | --- | --- | --- | --- | --- | --- | --- | --- | --- |
|  | R | D | U | T | R | D | U | T | R | D | U | T | R | D | U | T |
| *M. ochraceiventer*<br>(N=2) | 0 | - | - | 0 | 1<br>(50%) | - | - | 1<br>(50%) | 2<br>(100%) | - | - | 2<br>(100%) | 1<br>(50%) | - | - | 1<br>(50%) |
| *M. whiteheadi*<br>(N=12) | 1<br>(9%) | 0 | - | 1<br>(8%) | 0 | 0 | - | 0 | 11<br>(100%) | 1<br>(100%) | - | 12<br>(100%) | 0 | 0 | - | 0 |
| *N. cremoriventer*<br>(N=14) | 0 | 0 | - | 0 | 2<br>(20%) | 0 | - | 2<br>(14%) | 8<br>(80%) | 4<br>(100%) | - | 12<br>(86%) | 3<br>(30%) | 0 | - | 3<br>(21%) |
| *N. spp.*<br>(N=1) | 0 | - | - | 0 | 0 | - | - | 0 | 1<br>(100%) | - | - | 1<br>(100%) | 0 | - | - | 0 |
| *R. exulans*<br>(N=3) | 0 | 0 | - | 0 | 0 | 0 | - | 0 | 1<br>(50%) | 0 | - | 1<br>(33%) | 0 | 0 | - | 0 |
| *R. rattus* R3<br>(N=369) | 0 | 0 | 6<br>(4%) | 6<br>(2%) | 14<br>(16%) | 8<br>(6%) | 12<br>(9%) | 34<br>(9%) | 74<br>(85%) | 95<br>(70%) | 108<br>(77%) | 277<br>(75%) | 17<br>(20%) | 4<br>(3%) | 6<br>(4%) | 27<br>(7%) |
| *R. rattus tanezumi*<br>(N=41) | 11 | 0 | 0 | 0 | 2<br>(18%) | 0 | 11<br>(52%) | 13<br>(32%) | 10<br>(91%) | 6<br>(67%) | 15<br>(71%) | 31<br>(76%) | 1<br>(9%) | 1<br>(11%) | 4<br>(19%) | 6<br>(15%) |
| *R. tiomanicus*<br>(N=46) | 0 | 0 | 0 | 0 | 0 | 3<br>(9%) | 0 | 3<br>(7%) | 4<br>(40%) | 26<br>(79%) | 2<br>(67%) | 32<br>(70%) | 0 | 8<br>(24%) | 0 | 8<br>(17%) |
| *S. muelleri*<br>(N=327) | 0 | 0 | 0 | 0 | 4<br>(4%) | 5<br>(5%) | 15<br>(11%) | 24<br>(7%) | 90<br>(99%) | 102<br>(99%) | 132<br>(99%) | 324<br>(99%) | 16<br>(18%) | 23<br>(22%) | 66<br>(50%) | 105<br>(32%) |

Due to the abundance of ticks and their association with important zoonotic pathogens, we further examined their distribution across the urban-rural gradient (Figure 5). We found a strong association between urban environments and the presence of *Amblyomma* ticks, whereas *Ixodes* ticks were never encountered on a rodent from an urban site (P < 0.05). *Haemaphysalis* ticks showed divergent patterns across the urban-rural gradient, depending on the host: *S. muelleri* individuals in urban environments were more commonly infested than individuals in either rural or developing locations, whereas rural *R. rattus* were much more likely to be infested than individuals from either developing or urban locations (P < 0.05) (Figure 5). Bipartite networks constructed to examine the interactions between rodents and ticks at each location across the transect revealed that each of the rodent-tick networks were significantly modular. Networks from the rural and developing locations were similar in structure, with four modules in each and similar degrees of connectedness (Table 4). In contrast, the urban rodent-tick network was less fragmented, with only two modules and a higher degree of connectedness across the network. Examination of the top 20 most central nodes (i.e., those with the highest eigenvalues) in the projected unipartite networks also revealed similarities between the rural and developing locations, with a large number of collection sites and rodent species contributing to tick transmission at each location (Table 5). In contrast, all 20 of the most central nodes in the urban network were occupied by individuals belonging to *S. muelleri*, which were collected from only five sites.

**Figure 5.**
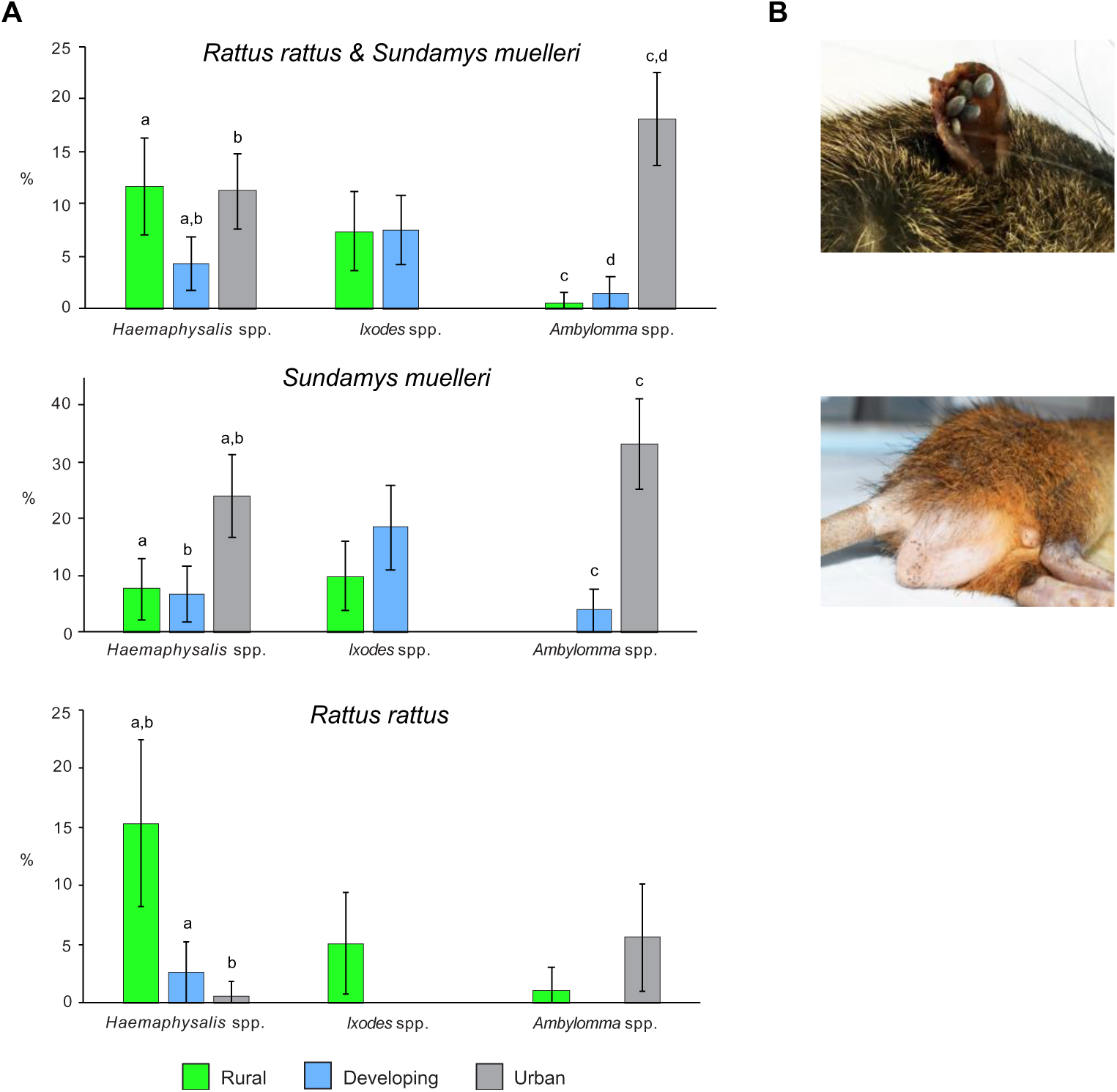
**(A) Proportion (%) of rodents with ticks identified as either *Haemaphysalis* spp., *Ixodes* spp., or *Amblyomma* spp. across the urban-rural gradient.** Individuals belonging to *S. muelleri* and *R. rattus* are shown both together and separately. Significant differences in the proportion of individuals with ticks at each location is indicated by lower case letters. **(B)** Photographs showing ticks *in situ* on *S. muelleri* from the urban location.

**Table 4.** Metrics associated with bipartite networks describing the interactions between rodents and ticks or microbes at each location across the urban-rural gradient. The location, connectedness (connect), modularity (number of modules), nestedness temperature (temp), and edge density (ED) with 95% confidence intervals are shown.

| Network | Location | Connect | Modularity<br>(N) | Temp | ED (95% CI) |
| --- | --- | --- | --- | --- | --- |
| Rodent:tick | Rural | 0.197 | 0.680 (4) | 18.73 | 0.32 (0.35-0.41) |
|  | Developing | 0.177 | 0.660 (4) | 25.94 | 0.25 (0.21-0.26) |
|  | Urban | 0.312 | 0.557 (2) | 19.48 | 0.52 (0.50-0.55) |
| Rodent:microbe | Rural | 0.145 | 0.711 (5) | 6.33 | 0.24 (0.21-0.29) |
|  | Developing | 0.115 | 0.586 (8) | 7.31 | 0.32 (0.29-0.36) |
|  | Urban | 0.107 | 0.511 (8) | 13.51 | 0.37 (0.29-0.51) |

**Table 5.**
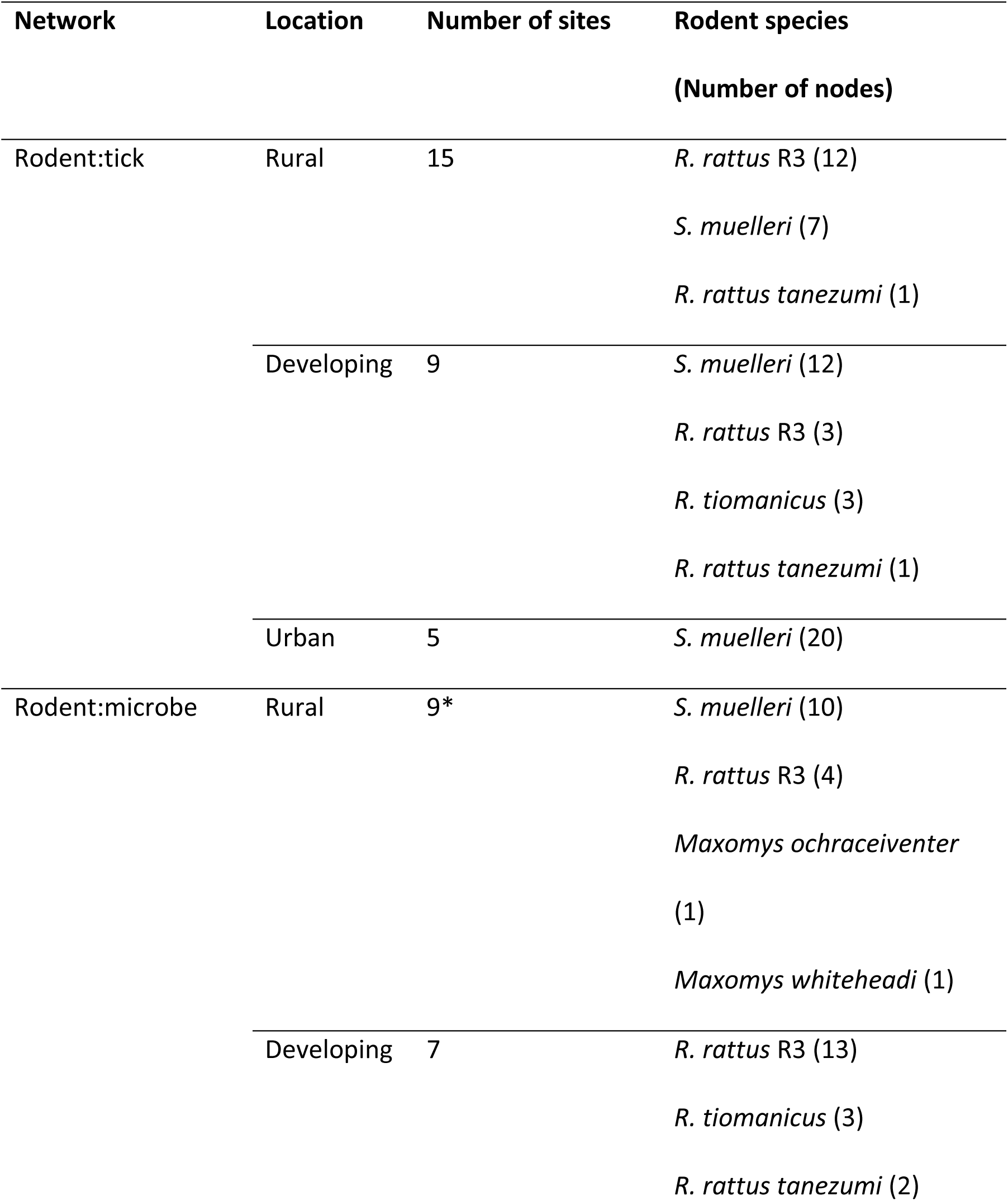
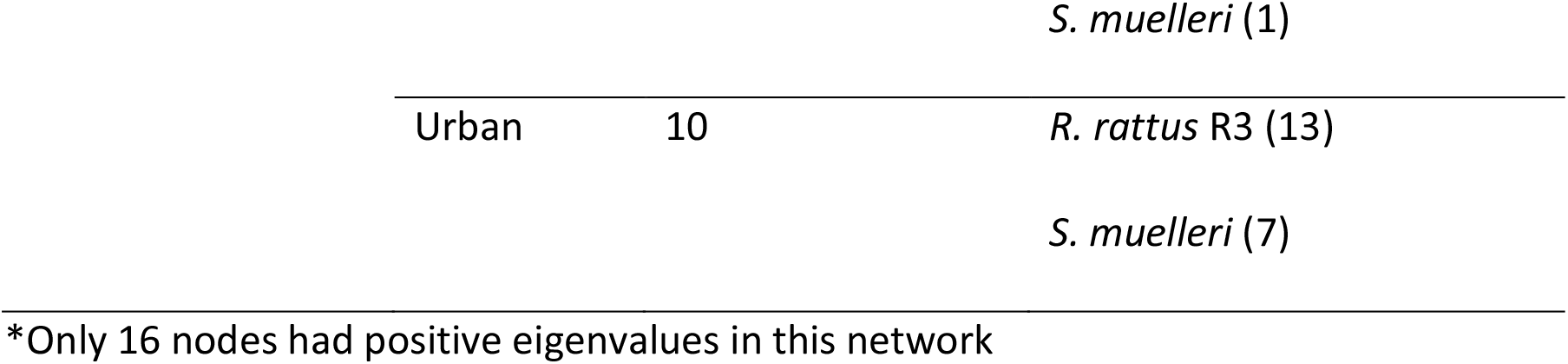
Rodent species and number of collection sites represented in the top 20 central nodes of interaction networks between rodents and their parasites, identified using Eigenvalue centrality. Eigenvalues for each node were calculated from unipartite rodent networks projected by connecting rodents based on tick or microbe sharing.

### Microbes and the urban-rural gradient

At least one of the 316 rodents tested yielded positive results by the assays used to detect mammarenaviruses (GenBank Accession #: MG999643), coronaviruses (MH107323-MH107330), orthohantaviruses (MH236804 - MH236810), orthohepeviruses (MH236817 - MH236830), paramyxoviruses (MH236811 - MH236816), and *T. gondii*, as well as *Bartonella* spp. (MG807665 - MG807845) and *Leptospira* spp. (sequences available at Blasdell et al. 2019b) (Supplemental Table 4). No animals were positive for alphaviruses, flaviviruses, enteroviruses, *Rickettsia* spp. or *Yersinia pestis* by the assays used in this study. While a single, previously described species was recovered from the mammarenavirus (*Wenzhou mammarenavirus*, WENV), orthohantavirus (Serang virus, SERV; a variant of *Thailand orthohantavirus*), orthohepevirus (*Orthohepevirus C* genotype 1, HEV-C1), and *T. gondii* PCR assays, multiple species or genotypes were detected by the remaining assays. Phylogenetic analysis of the coronavirus sequences revealed the presence of two clades of a single *Betacoronavirus* species that is most closely related to several *Betacoronavirus* lineage A isolates from Southeast Asian rodents (subgenus *Embecovirus*), as well as Human coronavirus HKU1 (Supplemental Figure 2). No clustering by host species was observed within this group. In contrast, phylogenetic analysis of the paramyxovirus sequences revealed two distinct species. The first was associated with *S*. *muelleri* (putatively named Rodent paramyxovirus SM) and appears distantly related to the *Morbillivirus* genus, along with viruses sequenced from *R. rattus* in Nepal (PREDICT_PMV-55/PR0225) and *Arvicanthis* sp. in Zambia (Rodent Paramyxovirus LR11-142). The second paramyxovirus (putatively named Rodent paramyxovirus R3) may represent a new species of *Jeilongvirus* (along with PREDICT_PMV-58/VN13M0007 from *Rattus* sp. in Viet Nam), and was detected in the kidneys of three *R. rattus* (lineage R3) (Supplemental Figure 3). Multiple species of the bacteria *Bartonella* and *Leptospira* were also identified and the results of these assays have been reported in detail previously (Blasdell et al. 2019a,b). However, *Bartonella* spp. was the most commonly detected pathogen in this study, with 57% of rodents infected across five species. Several known zoonoses were identified in the rodents sampled, including *B. elizabethae* and *B. tribocorum* (both identified in urban *R. rattus*), and *B. rattimassiliensis* (present in *R. rattus* across the entire urban-rural gradient). We also identified the two most significant causes of leptospirosis globally (*L. interrogans* and *L. borgpetersenii*), which were detected in 32% of rodents, combined. Overall, the prevalence of infection was low for most of the microbes surveyed here; however, the diversity of rodent-borne microbes identified in this study increased with increasing urbanization, and some pathogens (SERV, WENV), were only present at urban sites (Figure 6).

**Figure 6.**
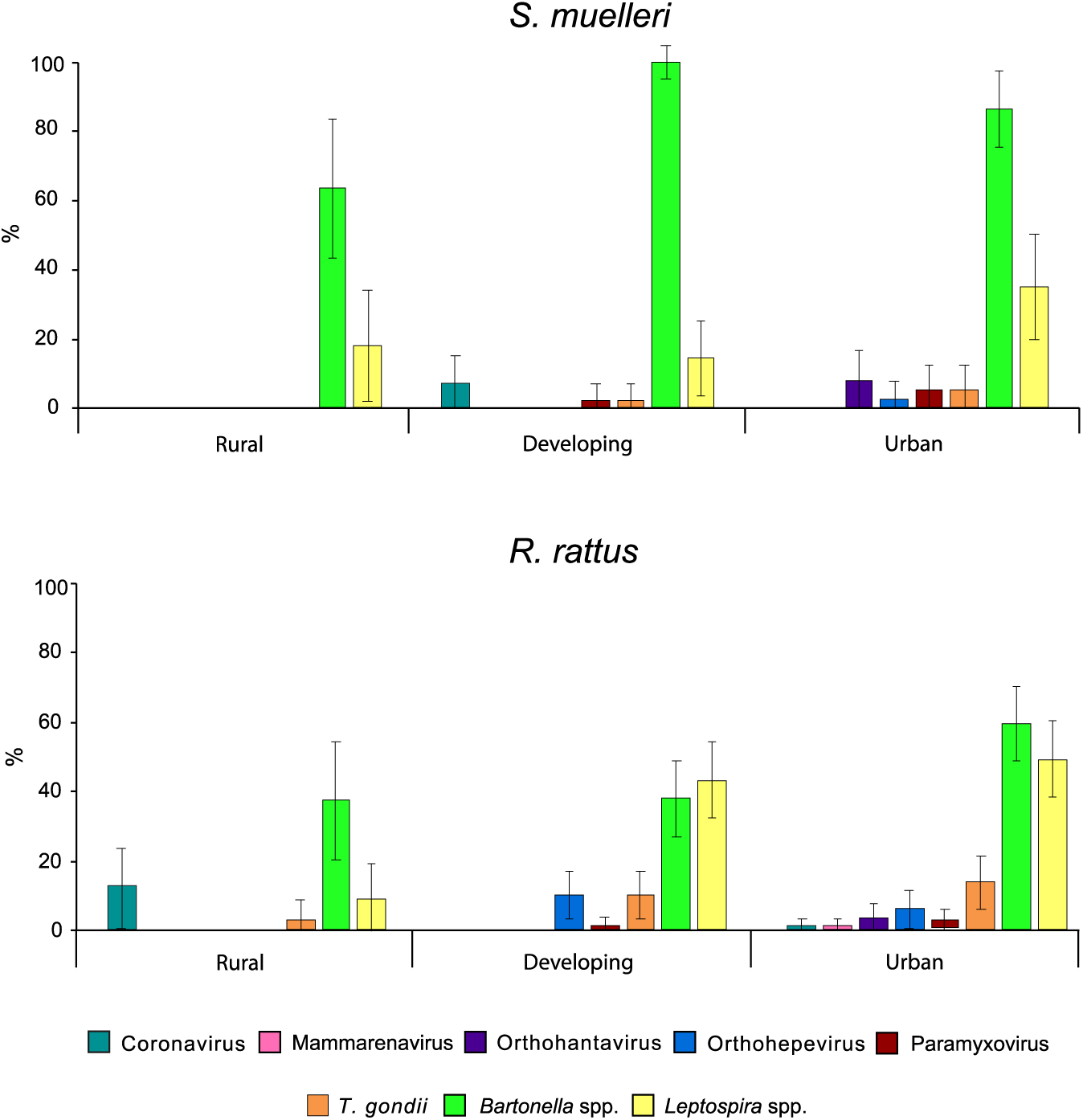
Proportion (%) of individuals belonging to *S. muelleri* and *R. rattus* infected by each pathogen at rural, developing, and urban locations across the urban-rural gradient. Pathogens were identified using consensus PCR assays and confirmed by Sanger sequencing.

An examination of the bipartite networks constructed to examine interactions between rodents and microbes across the urban-rural gradient revealed decreasing modularity with increasing anthropogenic influence in the landscape (Table 4). Networks from the developing and urban locations were similar in structure (eight modules each) indicating a higher degree of connectedness, whereas the rural-microbe network was more partitioned. Examination of the top 20 most central nodes in the projected unipartite networks also revealed similarities between the developing and urban locations that appear to be driven by the importance of *R. rattus* (65% of the top 20 central nodes were occupied by *R. rattus* in the developing and urban networks) (Table 5). In contrast, 63% of the most central nodes in the rural network were occupied by *S. muelleri*, with *R. rattus* occupying only one quarter of central nodes.

## Discussion

### Rodents and the urban-rural gradient

In this study, we found that rodent species diversity decreased with increasing anthropogenic influence on the landscape. No new species were found in the developing or urban locations that were not also present in the rural location, suggesting a steady loss of community complexity across the urban-rural gradient. This loss of species diversity has been observed repeatedly across taxa and is a predicted outcome of urbanization that appears to be driven by inter-species variation in traits associated with urban adaptability (Blair 1996, McKinney 2002, Croci et al. 2008, Grimm et al. 2008, Isaac et al. 2014, Santini et al. 2019). For wildlife to persist in cities, they must adapt to or avoid the risks associated with new environmental conditions (e.g., impervious surfaces, traffic, light pollution, etc.), while exploiting the novel resources and habitats created by urbanization (Grimm et al. 2008, Santini et al. 2019). For example, the species lost between the rural and developing locations in this study are considered lowland forest specialists (e.g., *M. rajah)*, while those lost between the developing and urban locations (e.g., *M. whiteheadi*) are associated with primary or secondary forests that are less frequent in urban areas (Wells et al. 2007, 2014, Charles & Ang 2010, Ruedas 2016). It may be that a strong preference for forested habitats is associated with specialized diets and reduced behavioral plasticity in some species that limits their ability to adapt to a high-risk, high-reward urban lifestyle (Santini et al. 2019). Indeed, only three species were found in habitats across the entire urban-rural gradient: *R. rattus* (lineages R3 and *tanezumi*), *R. tiomanicus*, and *S. muelleri*. Two of these, *R. rattus* (an introduced species) and *S. muelleri* (a native species) were found consistently across sites and were the most common species trapped in this study. These species exhibit many of the traits previously associated with successful urban rodents, including large body mass, large litter sizes, and generalist diets (Santini et al. 2019).

*Rattus rattus* is an urban exploiter and human commensal that is considered a highly invasive pest species globally. As an opportunistic omnivore with habitat preferences that range from forests to urban areas, *R. rattus* has frequently been cited as contributing to the local extinction of other wildlife (Towns et al. 2006, Stokes et al. 2009, Shields et al. 2014, Wells et al. 2014, Cusack et al. 2015). Critically, this species is also a known reservoir of a range of important zoonotic pathogens, including *Leptospira* spp., *Bartonella* spp., and hepatitis E virus (Meerburg et al. 2009, Himsworth et al. 2013, Mulyanto et al. 2013, Strand & Lundkvist 2019). In this study, we found that *R. rattus* was significantly more likely to be present at urban sites than developing or rural sites, and strongly preferred those with human infrastructure.

Consequently, we found that rural animals had significantly lower body mass indices than those trapped in locations with higher anthropogenic influence, which may be the result of decreased resource provisioning or reduced lifespan in rural areas. This is consistent with the hypothesis that *R. rattus* benefits from a close association with people and the accompanying resources and infrastructure we provide (Banks and Smith 2015). Although *R. rattus* was strongly associated with the urban location in this study, we found that classifying sites into ‘rural’, ‘developing’, or ‘urban’ categories alone was not a good predictor of the presence of *R. rattus* (or *S. muelleri*) across the landscape. This suggests that features of the local environment (i.e., at site level) may be more important predictors of species diversity and abundance than landscape context. Indeed, we found evidence that re-classifying our sites using nine fine-scale environmental variables better predicted species presence than location alone, especially for *R. rattus* (Figure 3). Specifically, sites with sewers, rubbish, buildings, and roads were highly correlated with the presence of *R. rattus* across the urban-rural gradient, while extensive green space and rivers were associated with an absence of this species.

In contrast, *S. muelleri* was most prevalent in the developing location, which may suggest a preference for disturbed or fragmented landscapes that provide access to both natural habitat and the increased resources associated with human habitation. Although *S. muelleri* is native to the lowland forests of Southeast Asia, individuals have been observed occupying patches of remnant vegetation, parks, and fallow land in urban and developing areas of Borneo (Madinah et al. 2013, Wells et al. 2014, Ng et al. 2017). *S. muelleri* has also been found to favor disturbed habitats and to achieve larger population sizes in habitat fragments due to a loss of predators from these environments (Charles 1996, Charles & Ang 2006, 2009). Together, these data strongly suggest that *S. muelleri* should be considered an urban adaptor species in Southeast Asia. We were less successful in predicting the presence of *S. muelleri* than *R. rattus* based on fine-scale environmental variables, due to a smaller number of individuals captured and a lower proportion of sites with *S. muelleri* present. However, we did find evidence to suggest that green sites adjacent to rivers and with an absence of domestic agriculture were more likely to support *S. muelleri*, which is consistent with previous suggestions that S*. muelleri* is a riverine species (Charles & Ang 2009). Surprisingly, the presence of roads was also associated with *S. muelleri* in our analysis. Roads occur along with many forms of both active (e.g., homes and agriculture), and inactive land use (e.g., fallow lands). As *S. muelleri* has previously been found at high densities in fallow lands, the association we detected between roads and *S. muelleri* may reflect a preference for disturbed habitats, for which roads are likely to be a proxy (Wells et al. 2014). While we were unable to explicitly account for the effects of interspecific interactions on the distribution of rodents in this study due to sampling limitations, we did observe an increase in the proportion of sites occupied by both *R. rattus* and *S. muelleri* with increasing urbanization across the gradient. Previous studies have demonstrated that the presence of invasive *R. rattus* can have both direct (e.g., interference through aggressive encounters) and indirect (e.g., decreased resource availability) impacts on the viability of native species in shared habitat patches (Harris and MacDonald 2007, Stokes et al. 2009). However, the majority of these studies were performed in natural environments and it is possible that competitive exclusion may explain the lower levels of co-occurrence we observed in the rural location. We hypothesize that the increased resources available in urban areas, coupled with the larger body size and aggressive nature of *S. muelleri*, allows this urban adapter species to co-exist with *R. rattus* when suitable habitat is limited.

### Ectoparasites and the urban-rural gradient

Consistent with previous studies, mesostigmatid mites were the most common ectoparasite carried by all rodents in this study, followed by ticks, lice, and fleas (Ng et al. 2017, Paramasvaran et al. 2009). *S. muelleri,* in particular, was relatively highly parasitized – nearly 100% of individuals were infested with mites and a third with ticks. Previous research on the ectoparasites carried by rodents in Malaysia have reported both low and high levels of ectoparasite diversity and abundance on the same rodent species studied here, across a range of habitats (Madinah et al. 2013, 2014, Ng et al. 2017). This variability was also observed in our data, as the proportion of individuals with ectoparasites varied by location and host species. This suggests that both the environment and the composition of the rodent community can influence the diversity and abundance of ectoparasites across a landscape.

In this study, we explored the local ecology of rodent-associated ticks in more detail to examine the impacts of urbanization on the abundance and distribution of ectoparasites. We identified three genera of hard ticks in this study: *Amblyomma*, *Haemaphysalis*, and *Ixodes*, each of which are considered to be amongst the most medically-important ectoparasites worldwide (Pfaffle et al. 2013). Interestingly, the abundance and distribution of each genus varied significantly across the urban-rural gradient (Figure 5). While the prevalence of *Amblyomma* ticks was significantly higher in the urban location, the distribution of *Ixodes* spp. displayed the opposite pattern – no individuals from the urban location were parasitized by *Ixodes* ticks, whereas *Haemaphysalis* spp. were found across the urban-rural gradient. Urbanization can impact the abundance and distribution of ticks by altering the availability of suitable hosts or by changing the external environment used by ticks when off-host, where they may spend up to 99% of their lifespan (McCoy et al. 2013, Pfaffle et al. 2013). Ticks have limited independent dispersal abilities, and as a result, their presence at a site is primarily influenced by where they detach from their hosts once engorged. For a population of ticks to be maintained at a site over time, suitable hosts for all life stages must be present in an environment that also supports off-host behaviors like questing (McCoy et al. 2013, Pfaffle et al. 2013, Estrada-Pena and de la Fuente, 2014).

In this study we found that rodents, and *S. muelleri* in particular, appear to be suitable hosts for both the larval and nymph stages of all three genera of ticks across the landscape. In contrast, adult ticks were found relatively infrequently, particularly in the urban location. This is consistent with the lifecycle of most hard-bodied ticks, where a single blood meal is taken over several days from a small vertebrate host during each of the larval and nymph stages, and a larger-bodied vertebrate (e.g., a large carnivore or ungulate) is preferentially fed on by adults (Estrada-Pena and de la Fuente, 2014). Although host preferences are determined by a range of factors, including host activity level, nesting behavior, grooming, and immune response, when a preferred host is not available, hungry ticks will attach to a sub-optimal host (Cumming 2004, van Duijvendijk et al. 2015). Thus, our observation that *S. muelleri* was more likely to host all three genera and life stages of ticks than *R. rattus* across the urban-rural gradient does not necessarily indicate that *S. muelleri* is the preferred host. Instead, it may suggest that in the context of this urbanizing landscape, *S. muelleri* is more likely to be present in the preferred microhabitats of these ticks. For example, individual *S. muelleri* were most heavily parasitized in the urban location, where they were restricted to green patches (no individuals were trapped within the built environment at any site), and observed at higher densities, particularly in comparison to rural sites. *S. muelleri* was also frequently the only small mammal captured at any urban site with a significant proportion of green space, which may be driving the importance of this species in the highly connected rodent:tick interaction networks constructed for the urban location. Therefore, we hypothesize that the dense populations of *S. muelleri* found at vegetated urban sites are favorable to the maintenance and transmission of *Amblyomma* and *Haemaphysalis* ticks in the urban environment. In contrast, the complete absence of *Ixodes* spp. from across the urban location suggests that they may have specific habitat or host requirements not present at our urban sites (Cumming 2004). Interactions between the local environment and the composition of the rodent community may also explain the increased tick parasitism of *R. rattus* in the rural location, as contact between *R. rattus* and vegetation suitable for questing is much more likely to occur in rural environments. Unfortunately, our ability to identify key features in the environment that may support tick presence was limited by the relatively small number of individuals infested with ticks in our data set and our inability to identify ticks to the species level.

### Microbes and the urban-rural gradient

Overall, we found the abundance and diversity of microbes carried by rodents increased with increasing urbanization across the gradient. Of the eight groups of microbes detected by consensus PCR, all were identified in the urban setting. In contrast, only *Bartonella* spp., *Leptospira* spp., *T. gondii,* and the betacoronaviruses were identified in rural animals. Notably, the prevalence of all viruses detected in this study was low. However, SERV, WENV, and HEV-C1 are known zoonotic pathogens, while the *Embecovirus* subgenus of betacoronaviruses contains several viruses capable of infecting people, including HCoV-OC43 and HCoV-HKU1 (Pattamadilok et al. 2006, Jonsson et al. 2010, Gamage et al. 2011, Decaro & Lorusso 2020, Reuter et al. 2020).

Two pathogens were found only in the urban location – SERV and WENV. Rodents are the primary reservoir hosts for both orthohantaviruses and mammarenaviruses, and transmission between animals occurs through both vertical and horizontal routes (Banerjee et al. 2008. Calisher et al. 2009). Human infection with these viruses most commonly occurs through the inhalation of aerosolized infectious excreta, although transmission through rodent bites has also been documented (Schultze et al. 2002). In this study, SERV was detected in *R. rattus* from a single site (a large market in the city center – also the only site where WENV was detected), and in *S. muelleri* from three vegetated sites, two of which also contained *R. rattus.* It is not clear why these rodent-borne viruses were not detected in the developing or rural locations; however, several factors may explain their absence. These include: (i) lower population density and/or smaller population size of competent hosts in the rural and developing locations, which may reduce transmission below the level necessary for virus persistence; (ii) the presence of less-competent hosts in these more diverse rodent communities, which may reduce intraspecific interactions and horizontal transmission (i.e., the dilution effect); (iii) recent introductions of infected rodents through shipping or other routes of importation into Kuching; or (iv) an artefact of our relatively small sample size (both in number of animals and number of sites) that may have inadvertently excluded microhabitats more conducive to viral transmission in these locations (Clay et al. 2009, Dobly 2012, Voutilainen et al. 2012, Raharinosy et al. 2018). It is therefore possible that these (and other) zoonotic pathogens are present in rodents at low frequency across the landscape, yet were not detected here. Notably, both SERV and WENV have been recently described in Southeast Asia and little is known about the epidemiology or public health risks associated with these viruses (Plyusnina et al. 2009, Blasdell et al. 2016). However, the presence of SERV at multiple urban sites with differing microhabitats, and in at least two reservoir species, suggests that this virus is endemic to the local area and may pose a risk to people where contact with rodents is high. In contrast, our singular detection of WENV from *R. rattus* in Kuching’s city center suggests that the risk of zoonotic transmission of this virus is likely minimal. However, as many rodent-borne pathogens have a similar clinical presentation and are frequently mis-diagnosed, the public health significance of these viruses may be underestimated (Pattamadilok et al. 2006, Meerburg et al. 2009, Gamage et al. 2011, Chen et al. 2014, Blasdell et al. 2016). As a result, targeted human serosurveys will be necessary to determine if the presence of the zoonotic viruses identified in urban rodents in this study are associated with health risks to people.

We found evidence of HEV-C1 (rat origin hepevirus) and two host-associated paramyxoviruses in rodents from both the developing and urban locations in this study. HEV-C1 was predominantly identified from *R. rattus* lineage R3 (86% of detections) at sites with high rodent density, a high proportion of built infrastructure, and substantial resource provisioning from adjacent markets and restaurants. HEV-C1 has previously been detected in a wide range of rodent species across three continents (Asia, Europe, North America), and occasional spillovers into other mammals, including people, have been documented (Andonov et al. 2019, Pallerla et al. 2020, Reuter et al. 2020, Sridhar et al. 2018, 2020). Globally, zoonotic transmission of HEV-C1 through contact with rodent feces appears infrequent. However, there is a paucity of data as commercially available tests for HEV are designed to detect *Orthohepevirus A* (human HEV) only, and these have failed to detect clinical HEV-C1 infections in the past (Sridhar et al. 2018, 2020, Andonov et al. 2019, Reuter et al. 2020). Similarly, although a growing number of rodent-borne paramyxoviruses have been sequenced in the past decade, little is known about the zoonotic potential of these viruses. We found two distinct species of paramyxovirus in this study - Rodent paramyxovirus SM was identified in the kidneys of three *S. muelleri* from two urban and one developing site, whilst Rodent paramyxovirus R3 was detected in the kidneys of three *R. rattus* from one urban and one developing site. Rodents are second only to bats as the most common wildlife hosts of paramyxoviruses, yet there have been no confirmed instances of zoonotic transmission of rodent-borne viruses. In contrast, several bat-borne paramyxoviruses have spilled over into the human population in recent years, with significant public health impacts (Drexler et al. 2012, Onyuok et al. 2019). This is perhaps unexpected, as most rodent-borne paramyxoviruses have been associated with the kidney and could potentially be transmitted through contact with infected urine, mimicking routes of exposure for bat-borne paramyxoviruses such as Nipah virus (Drexler et al. 2012).

In this study, only *Bartonella* spp., *Leptospira* spp., and *T. gondii* were regularly identified in rodents from all three locations across the urban-rural gradient, and in each case, prevalence was highest in the urban location. All of these are well-known rodent-borne pathogens with significant zoonotic potential, and are also most commonly transmitted between animals via the environment (*Leptospira* spp., and *T. gondii*), or an ectoparasite vector (*Bartonella* spp.) (Saisongkorh et al. 2009, Costa et al. 2015, Regier et al. 2016). We found that *R. rattus* was most commonly associated with the presence of all three species of *Bartonella* that are known to be zoonotic (*B. elizabethae, B. tribocorum, B. rattimassiliensis*); however, urban *S. muelleri* were more likely to be infected with any *Bartonella* species (Blasdell et al. 2019a). We found that high densities of *S. muelleri* that were frequently observed in patches of remnant vegetation, parks, and fallow land across the urban location were associated with both a high ectoparasite burden (e.g., *Haemaphysalis* ticks), and a high prevalence of *Bartonella* spp.. These habitats are frequently used by people in and around Kuching for recreational and foraging activities, which suggests potential local ‘hot-spots’ for zoonotic transmission. To our knowledge, only ocular bartonellosis has been reported in Malaysia, which is associated with *B. henselea*, a species that was not identified here (Blasdell et al. 2019a, Tey et al. 2020). However, as with many other rodent-borne diseases, *Bartonella* infections generally result in undifferentiated febrile illnesses that are often misdiagnosed, and as a result, the public health impacts of this pathogen are not well understood (Saisongkorh et al. 2009, Regier et al. 2016).

We found that all *Leptospira* (and *L. borgpetersenii* in particular) were statistically more likely to be present at sites with higher anthropogenic activity, as indicated by the presence of both commercial and residential activity at a site (Blasdell et al. 2019b). This finding was unexpected, as leptospirosis is predominantly considered a rural disease and recent Malaysian outbreaks have been associated with rural recreational activities (Shafei et al. 2012, Costa et al. 2015, Benacer et al. 2016). However, infection prevalence has been found to be higher in wildlife occupying urban habitats, and this trend appears to be particularly significant for rodents (Villanueva et al. 2010, Andersen-Ranberg et al. 2016, Benacer et al. 2013). Notably, the human commensal, *R. rattus*, was most commonly infected in this study. *R. rattus* is able to reach high population densities in urban areas, and is commonly found within the built environment, where contact with people is more likely to occur (Feng and Himsworth 2014). This suggests that the high prevalence of *Leptospira* spp. in urban *R. rattus* found here may be associated with a significant risk of human infection in this area. *R. rattus* was also the primary host of *T. gondii* amongst sampled rodents, as only three other individuals (all *S. muelleri*) were positive at any site. *T. gondii* was the only pathogen in this study whose prevalence increased consistently with increasing urbanization across the gradient, and this trend may be driven by the importance of domestic cats as the definitive hosts of *T. gondii*. Cats were strongly associated with human habitation across our study sites and were much more frequent in the urban location (Hong & Mohd-Azlan 2018). Although rodents play a key role in the transmission cycle of *T. gondii* by acting as a primary source of infection for cats, people (and other mammals) become infected through the ingestion of food, water, or soil contaminated with cat feces and the oocysts they contain, which can remain infectious for up to a year in humid conditions (Dubey & Frenkel 1998, Dubey 2010, Robert-Gangneux and Darde 2012). This may help explain the association between *R. rattus* and *T. gondii* in the built environment, where *R. rattus* was frequently captured in and around sewer systems that are conducive to water-borne transmission (Gotteland et al. 2014). Although there has been little research on the biological or ecological factors that influence *T. gondii* prevalence in cities, the local environment in Kuching appears to contain several risk factors for human infection. These include high average rainfalls, open sewers, large numbers of cats, and an infection prevalence in *R. rattus* at the higher end of published reports (Robert-Gangneux and Darde 2012, Gotteland et al. 2014). Human infections with *T. gondii* appears to be frequent in Malaysia, with a seroprevalence in healthy people that ranges from 14-30% and varies with ethnicity, sex, age and socioeconomic status (Nissapatorn & Abdullah 2004, Nissapatorn 2010). This suggests that zoonotic transmission of *T. gondii* is relatively frequent in this area, and may be driven by the increased prevalence of the parasite in areas with high densities of cats, rodents, and people in an environment well-suited to zoonotic transmission.

### Urbanization and human disease risk

In this study, we explored how the abundance and distribution of rodents and their ectoparasites is influenced by increasing urbanization across a landscape. The goal of this work was to begin to identify habitats, species, or behaviors that may alter the risk of zoonotic disease emergence in tropical cities. Predictably, we found that rodent species diversity decreased with increasing urbanization, leaving an urban species assemblage that was dominated by two species – *R. rattus* (an urban exploiter) and *S. muelleri* (an urban adapter). Although these species were occasionally found in the same microhabitats across the urban-rural gradient, habitat partitioning was more common – *R. rattus* was most frequently found within the built environment, while *S. muelleri* was associated with green patches. Spatial segregation between human commensals and native species has been observed previously and may be a common feature of cities (Baker et al., 2003; Castillo et al., 2003; Parsons et al., 2018). As a result, the location, frequency, and intensity of contact between people and urban exploiters/adapters is likely to differ, as are the implications for zoonotic disease risk.

Like many urban adapter species, *S. muelleri* demonstrated a clear preference for vegetated sites in urban areas, where they were both abundant and heavily infested with *Haemaphysalis* and *Amblyomma* ticks (Baker et al. 2003, Cavia et al. 2009). Ticks are capable of transmitting a greater diversity of pathogens than other ectoparasites, and both *Haemaphysalis* and *Amblyomma* ticks have been associated with zoonotic pathogens, including *Rickettsia* spp., *Bartonella* spp., *Borrelia* spp., and tick-borne encephalitis virus (Parola & Raoult 2001). Ticks can become infected with a pathogen during blood feeding at any life stage; however, ticks that feed on the same host species across multiple life stages are more likely to become infected and to transmit this infection during subsequent feedings (Mihalca and Sandor 2013, van Duijvendijk et al. 2015). In this context, the dense urban populations of *S. muelleri* that were heavily infested with multiple life stages of ticks may pose a particular risk to people should a zoonotic pathogen be present in the population. As in many Southeast Asian cities, green spaces across Kuching are heavily utilized by residents for recreational and other activities, and are often considered to be clean, healthy urban environments. However, the results of this study suggest that these activities may pose under-appreciated public health risks akin to those associated with the transmission of Lyme disease in temperate urban green spaces (Estrada-Pena and de la Fuente 2014, Blasdell et al. 2019a). We suggest that effective public health messaging could be used to inform residents of tropical cities about the potential risks posed by ticks in urban green spaces, and encourage the use of personal insect repellants and other preventative measures. These efforts should be supplemented with a targeted pest management plan to reduce rodent density in areas of high abundance and where frequent contact with people is likely.

As has been found for other human commensals, including *R. norvegicus* and *M. musculus*, the presence of *R. rattus* was strongly correlated with the presence of built infrastructure in the urban environment (Feng & Himsworth 2014, Himsworth et al. 2013). Although this association likely translates to a reduced risk from tick-borne zoonoses, our results also suggest that there may be an elevated risk from multi-host pathogens capable of environmental transmission (e.g., *Leptospira* spp, *T. gondii*, HEV-C1) in habitats where *R. rattus,* and other commensals, thrive. This suggests that some features of the built environment may support pathogen persistence, including open sewers, wet markets, and co-located commercial and residential activity (Blasdell et al. 2019b, Strand & Lundkvist 2019, Hagan et al. 2016). Further studies on the relationships between features of the built environment, the presence of pathogens, and the frequency of zoonotic transmission at a fine scale will be needed to further understand how the structure and use of cities may influence zoonotic disease risk.

In sum, we propose that impact of urbanization on zoonotic disease risk is a result of interactions between the response of reservoir hosts, ectoparasite vectors, pathogens, the surrounding environment, and human behavior (Figure 7). The process of urbanization can act to change zoonotic disease risk by: (i) changing the abundance and distribution of available hosts and/or vectors, thereby impacting pathogen persistence; (ii) changing the community composition of hosts/vectors, altering inter- and intra-specific interactions as well as host/vector competence; (iii) changing the behavior, movement patterns, and immune status of hosts, thereby altering disease susceptibility and transmission; (iv) changing the structure and utility of the environment in a manner that influences transmission (e.g., increased flooding, poor sanitation, high-density housing, etc.); and (v) changing the distribution or transmission dynamics of a pathogen directly. How these factors interact to influence human disease risk in cities will vary by geographic location, and the scale and speed of land-use change, among other factors. Given the rapid rate of urbanization in Southeast Asia and other tropical regions of the world, it will be critical to focus research efforts on cities in the tropics, where the public health consequences of a new or re-emerging infectious disease may have significant consequences.

**Figure 7.**
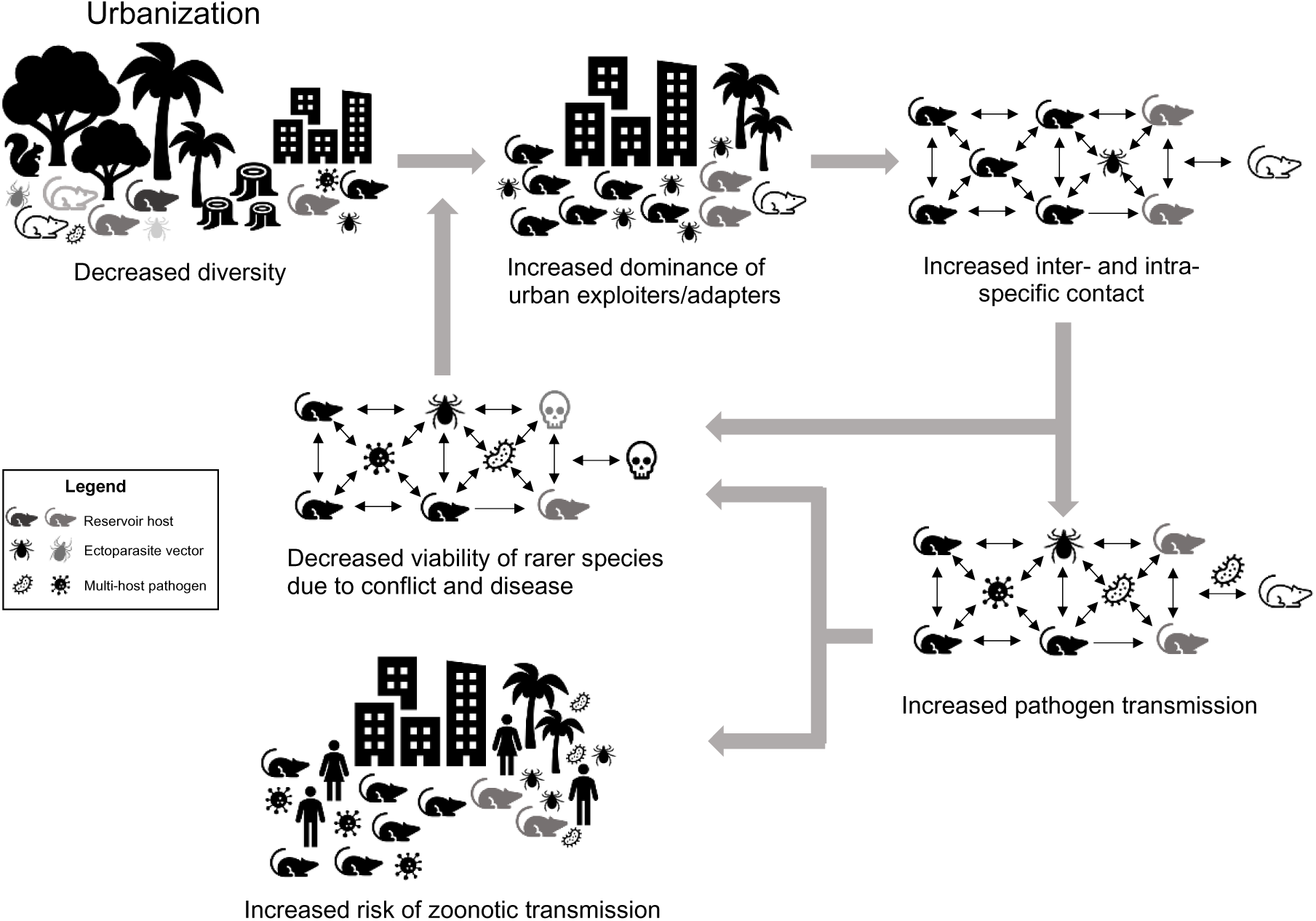
Proposed model of the impacts of urbanization on the transmission of multi-host pathogens and zoonotic disease risk. Urbanization results in decreased host diversity in cities and habitat partitioning of the remaining species. This leads to increased dominance of urban exploiters (solid black rodents), and urban adapters (solid gray rodents) in urban environments, as well as the ectoparasites that favor these hosts/habitats (black ticks). As inter- and intra-specific contact between urban exploiters, urban adapters, and their ectoparasites increases, so does multi-host pathogen transmission, which can lead to reduced viability of rarer species due to competition, predation, and disease. Together, these factors contribute to a heterogeneous distribution of hosts, vectors, and pathogens in the urban environment, with a higher risk of zoonotic disease transmission arising in habitats where frequent contact between hosts, vectors, pathogens and people are most likely to occur.

Our study had several significant limitations, the most important of which is the absence of data from people who represent the ‘end’ of the zoonotic transmission cycle. Although this study is one of the first to simultaneously examine the influence of urbanization on reservoir hosts, ectoparasites, and potential pathogens, the lack of relevant epidemiological, serological, or molecular data from people within our study area limits our ability to identify factors associated with genuine zoonotic transmission. Instead, we were only able to identify aspects of urbanization and the urban environment that may be considered risky given the presence of potential drivers of zoonotic infection in areas of high human activity. Furthermore, we were able to screen only a subset of the rodents collected in this study (and none of the ectoparasites), for potential zoonoses. A key next step will be to associate pathogens circulating in rodents, their ectoparasites, and people in the same locations. Similarly, by using a consensus PCR approach to microbial detection we may have missed important pathogens already circulating in this area (e.g., *Borrelia* spp.), that may have revealed the presence of other important drivers in this system (Lau et al. 2020). However, a metagenomics-based study of all microbes carried by these same rodents and their ectoparasites is currently underway and is expected to reveal a greater diversity of potential zoonoses across the urban-rural gradient (Harvey et al. 2019, Wu et al. 2021).

## Acknowledgments

The authors thank Andrew Alek Tuen (Universiti Malaysia Sarawak), Andrew Joris Noyen (Padawan Municipal Council), Basheer Ahmed (Kuching North Municipal Council), Danielle Levesque (Universiti Malaysia Sarawak), Dilop Jina (Padawan Municipal Council), Jean-Bernard Duchemin (CSIRO), Lee Trinidad (CSIRO), Nurshilawati Abdul Latip (Universiti Malaysia Sarawak), Patrick Lai Ganum (Kuching South Municipal Council), Rachel Amos-Ritchie (CSIRO), and Samuel Wong (Universiti Malaysia Sarawak).

