## Supplemental data for "Rats in the city: implications for zoonotic disease risk in an urbanizing world"

### SUPPLEMENTAL MATERIAL – METHODS

#### Details of PCR-based microbial screening

**Alphavirus** (tissues: kidney, liver, spleen)

Sánchez-Seco MP, Rosario D, Quiroz E, Guzmán G, Tenorio A. A generic nested-RT-PCR followed by sequencing for detection and identification of members of the alphavirus genus. J Virol Methods. 200; 95:153-61.

**Coronaviruses** (tissues: kidney, lung)

Watanabe S, Masangkay JS, Nagata N, Morikawa S, Mizutani T, Fukushi S, Alviola P, Omatsu T, Ueda N, Iha K, Taniguchi S, Fujii H, Tsuda S, Endoh M, Kato K, Tohya Y, Kyuwa S, Yoshikawa Y, Akashi H. Bat coronaviruses and experimental infection of bats, the Philippines. Emerg Infect Dis. 2010;16:1217-23.

**Enteroviruses** (tissues: lung, feces)

Nix WA, Oberste SM, Pallansch MA. Sensitive, seminested PCR amplification of VP1 sequences for direct identification of all enterovirus serotypes from original clinical specimens. J Clin Microbiol. 2006;44:2698-2704.

Ly N, Tokarz R, Mishra N, Sameroff S, Jain K, Rachmat A, An US, Newell S, Harrison DJ, Lipkin WI. Multiplex PCR analysis of clusters of unexplained viral respiratory tract infection in Cambodia. Virol J. 2014;11:224.

**Flaviviruses** (tissues: spleen, liver)

Moureau G, Temmam S, Gonzalez JP, Charrel RN, Grard G, de Lamballerie X. A real-time RT-PCR method for the universal detection and identification of flaviviruses. Vector Borne Zoonotic Dis. 2007;7:467-477.

**Hantavirus** (tissues: kidney, lung)

Anthony SJ, Epstein JH, Murray KA, Navarrete-Macias I, Zambrana-Torrelio CM, Solovyov A, Ojeda-Flores R, Arrigo NC, Islam A, Ali Khan S, Hosseini P, Bogich TL, Olival KJ, Sanchez-Leon MD, Karesh WB, Goldstein T, Luby SP, Morse SS, Mazet JA, Daszak P, Lipkin WI. A strategy to estimate unknown viral diversity in mammals. mBio. 2013;4:e00598-13.

**Mammarenavirus** (tissues: lung, spleen)

Vieth S, Drosten C, Lenz O, Vincent M, Omilabu S, Hass M, Becker-Ziaja B, ter Meulen J, Nichol ST, Schmitz H, Günther S. RT-PCR assay for detection of Lassa virus and related Old World arenaviruses targeting the L gene. Trans R Soc Trop Med Hyg. 2007;101:1253-64.

**Orthohepevirus** (tissues: liver, spleen)

Firth C, Bhat M, Firth MA, Williams SH, Frye MJ, Simmonds P, Conte JM, Ng J, Garcia J, Bhuva NP, Lee B, Che X, Quan PL, Lipkin WI. Detection of zoonotic pathogens and characterization of novel viruses carried by commensal Rattus norvegicus in New York City. mBio. 2014;5:e01933-14.

**Paramyxovirus** (tissues: kidney, lung, spleen)

Tong S, Chern S-WW, Li Y, Pallansch MA, Anderson LJ. Sensitive and broadly reactive reverse transcription-PCR assays to detect novel paramyxoviruses. J Clin Microbiol. 2008;46:2652-8.

**Rickettsia spp.** (tissues: ear, kidney, liver, spleen)

Karpathy SE, Hayes EK, Williams AM, Hu R, Krueger L, Bennett S, Tilzer A, Velten RK, Kerr N, Moore W, Eremeeva ME. Detection of *Rickettsia felis* and *Rickettsia typhi* in an area of California endemic for murine typhus. Clin Microbiol Infect. 2009;15:218-9.

Roux V, Raoult D. Phylogenetic analysis of members of the genus Rickettsia using the gene encoding the outer-membrane protein rOmpB (ompB). Int J Syst Evol Microbiol. 2000;4:1449–55.

**Toxoplasma gondii** (tissues: lung)

Homan WL, Vercammen M, De Braekeleer J, Verschueren H. Identification of a 200- to 300-fold repetitive 529 bp DNA fragment in Toxoplasma gondii, and its use for diagnostic and quantitative PCR. Int J Parasitol. 2000;30:69-75.

**Yersina pestis** (tissues: liver, lung, spleen)

Stewart A, Satterfield B, Cohen M, O'Neill K, Robison R. A quadruplex real-time PCR assay for the detection of *Yersinia pestis* and its plasmids. J Med Microbiol. 2008;57:324-31.

### SUPPLEMENTAL MATERIAL - TABLES

**Supplemental Table 1. Site characteristics at each location**. The total number (%) of sites where each variable was recorded during rodent trapping.

| Category | Variable | **Rural** (N=65) | **Developing** (N=23) | **Urban** (N=27) |
|---|---|---|---|---|
| **Dominant land-use type** | Commercial | 0 | 0 | 3 (11%) |
| | Mixed Commercial/Residential | 0 | 4 (17%) | 10 (37%) |
| | Forest | 7 (11%) | 2 (9%) | 1 (4%) |
| | Scrub | 16 (25%) | 14 (61%) | 12 (44%) |
| | Village/Residential | 42 (65%) | 3 (13%) | 1 (4%) |
| **Next to or on ecotone*** | Commercial | 2 (3%) | 7 (30%) | 4 (15%) |
| | Village/Residential | 13 (20%) | 16 (70%) | 8 (30%) |
| | Scrub | 6 (9%) | 8 (35%) | 1 (4%) |
| | Forest | 48 (74%) | 1 (4%) | 1 (4%) |
| | Agriculture | 5 (8%) | 1 (4%) | 2 (7%) |
| | None | 9 (14%) | 6 (26%) | 15 (56%) |
| **Waterbody*** | Flowing (stream, river) | 34 (52%) | 10 (43%) | 9 (33%) |
| | Standing (swamp, pond) | 18 (28%) | 0 | 1 (4%) |
| | None | 20 (31%) | 13 (57%) | 17 (63%) |
| **Sewer** | Present | 24 (37%) | 14 (61%) | 23 (85%) |
| | Absent | 41 (63%) | 9 (39%) | 4 (15%) |
| **Livestock*** | Chickens | 50 (77%) | 7 (30%) | 5 (19%) |
| | Pigs | 10 (15%) | 0 | 0 |
| | None | 15 (23%) | 16 (70%) | 22 (81%) |
| **Food plants** | Yes | 38 (58%) | 5 (22%) | 4 (15%) |
| | No | 27 (42%) | 18 (78%) | 23 (85%) |
| **Rubbish** | Yes | 32 (49%) | 16 (70%) | 26 (96%) |
| | No | 33 (51%) | 7 (30%) | 1 (4%) |
| **Rubbish type*** | Old | 29 (45%) | 11 (48%) | 14 (52%) |
| | New | 9 (14%) | 9 (39%) | 14 (52%) |
| | N/A | 33 (51%) | 7 (30%) | 1 (4%) |
| **Roads** | Yes | 35 (54%) | 21 (91%) | 24 (89%) |
| | No | 30 (46%) | 2 (9%) | 3 (11%) |
| **Dominant building type** | Residential | 55 (85%) | 9 (39%) | 4 (15%) |
| | Mixed Commercial/Residential | 1 (2%) | 10 (43%) | 16 (59%) |
| | Commercial | 2 (3%) | 1 (4%) | 2 (7%) |
| | Industrial | 1 (2%) | 1 (4%) | 3 (11%) |
| | No building | 6 (9%) | 2 (9%) | 2 (7%) |
| **Commercial food storage/prep** | Yes | 11 (17%) | 7 (30%) | 9 (33%) |
| | No | 54 (83%) | 16 (70%) | 18 (67%) |
| **Average building † condition score** | 1 | 0 | 1 (4%) | 0 |
| | 2 | 4 (6%) | 2 (9%) | 1 (4%) |
| | 3 | 25 (38%) | 4 (17%) | 15 (56%) |
| | 4 | 30 (46%) | 12 (52%) | 9 (33%) |
| | 5 | 0 | 3 (13%) | 0 |
| **Dominant green space type** | Forest | 27 (42%) | 1 (4%) | 2 (7%) |
| | Scrub | 9 (14%) | 12 (52%) | 6 (22%) |
| | Garden | 8 (12%) | 1 (4%) | 5 (19%) |
| | Mixed | 21 (32%) | 7 (30%) | 3 (11%) |
| | None | 0 | 2 (9%) | 11 (41%) |

*More than one category may have been assigned to each site

† The condition of each building was scored as follows: Unused ruin=1, Poor=2, Fair=3, Good=4, Excellent=5; with the average value per site noted above.

**Supplemental Table 2. Number (N) of rodents caught by location across the urban-rural gradient**. For each sex (male (M), female (F), unknown (U), total (T), numbers are reported as N mature/N juvenile animals.

| Species | Rural (N mature/N juvenile) | | | | Developing (N mature/N juvenile) | | | | Urban (N mature/N juvenile) | | | | Total (N mature/N juvenile) | | | |
|---|---|---|---|---|---|---|---|---|---|---|---|---|---|---|---|---|
| | **M** | **F** | **U** | **T** | **M** | **F** | **U** | **T** | **M** | **F** | **U** | **T** | **M** | **F** | **U** | **T** |
| *R. rattus* R3* | 35/6 | 42/4 | 0/2 | 77/12 | 65/4 | 63/10 | 3/0 | 131/14 | 72/5 | 53/10 | 1/0 | 126/15 | 172/15 | 158/24 | 4/2 | 334/41 |
| *R. rattus tanezumi** | 4/0 | 6/1 | 0 | 10/1 | 3/1 | 4/1 | 0 | 7/2 | 13/1 | 5/2 | 0 | 18/3 | 20/2 | 15/4 | 0 | 35/6 |
| *R. tiomanicus** | 2/0 | 7/1 | 3/1 | 12/2 | 19/0 | 13/1 | 4/0 | 36/1 | 2/0 | 1/0 | 0 | 3/0 | 23/0 | 21/2 | 7/1 | 51/3 |
| *R. exulans* | 2/0 | 0 | 0 | 2/0 | 1/0 | 0 | 0 | 1/0 | 0 | 0 | 0 | 0 | 3/0 | 0 | 0 | 3/0 |
| *S. muelleri** | 43/3 | 43/2 | 0 | 86/5 | 51/7 | 43/2 | 3/0 | 97/9 | 55/12 | 62/4 | 1/0 | 118/16 | 149/22 | 148/8 | 4/0 | 301/30 |
| *N. cremoriventer* | 8/0 | 2/0 | 12/0 | 22/0 | 2/0 | 2/0 | 15/0 | 19/0 | 0 | 0 | 0 | 0 | 10/0 | 4/0 | 27/0 | 41/0 |
| *N.* spp. | 0 | 1/0 | 0 | 1/0 | 0 | 0 | 0 | 0 | 0 | 0 | 0 | 0 | 0 | 1/0 | 0 | 1/0 |
| *M. whiteheadi* | 6/0 | 4/1 | 0 | 10/1 | 0 | 0/1 | 0 | 0/1 | 0 | 0 | 0 | 0 | 6/0 | 4/2 | 0 | 10/2 |
| *M. ochraceiventer* | 1/0 | 1/0 | 2/0 | 4/0 | 0 | 0 | 0 | 0 | 0 | 0 | 0 | 0 | 0 | 0 | 0 | 0 |
| *M. rajah* | 0 | 0 | 1/0 | 1/0 | 0 | 0 | 0 | 0 | 0 | 0 | 0 | 0 | 0 | 0 | 1/0 | 1/0 |
| **Total** | 101/9 | 106/9 | 18/3 | 225/21 | 141/12 | 125/15 | 25/0 | 291/27 | 142/18 | 121/16 | 2/0 | 265/34 | 384/39 | 352/40 | 45/3 | 781/82 |

*Indicates species for which differences in trapping success by sex were assessed by an exact binomial test of goodness of fit ($P > 0.05$ in all cases).

**Supplemental Table 3. Life stages of ticks collected from 815 rodents in this study.** For each species, the number and proportion (%) of infested rodents are shown.

| **Species** | **Location (N sampled)** | ***Haemaphysalis* larvae** | ***Haemaphysalis* nymphs** | ***Ixodes* larvae** | ***Ixodes* nymphs** | ***Ixodes* adults** | ***Ambylomma* larvae** | ***Ambylomma* nymphs** | ***Ambylomma* adults** |
|---|---|---|---|---|---|---|---|---|---|
| *R. rattus* | Rural (N=98) | 9 (9.2%) | 7 (7.1%) | 1 (1.0%) | 0 | 4 (4.1%) | 1 (1.0%) | 0 | 0 |
| | Developing (N=151) | 3 (2.0%) | 1 (0.7%) | 0 | 0 | 0 | 0 | 0 | 0 |
| | Urban (N=161) | 0 | 1 (0.6%) | 0 | 0 | 0 | 6 (3.7%) | 3 (1.9%) | 1 (0.6%) |
| | Total (N=410) | 12 (2.9%) | 9 (2.2%) | 1 (0.2%) | 0 | 4 (1.0%) | 7 (1.7%) | 3 (0.7%) | 1 (0.2%) |
| *S. muelleri* | Rural (N=91) | 4 (4.4%) | 4 (4.4%) | 2 (2.2%) | 1 (1.1%) | 6 (6.6%) | 0 | 0 | 0 |
| | Developing (N=103) | 5 (4.9%) | 3 (2.9%) | 3 (2.9%) | 6 (5.8%) | 8 (7.8%) | 2 (1.9%) | 2 (1.9%) | 0 |
| | Urban (N=133) | 22 (16.5%) | 25 (18.8%) | 0 | 0 | 0 | 30 (22.6%) | 27 (20.3%) | 0 |
| | Total (N=327) | 31 (9.5%) | 32 (9.8%) | 5 (1.5%) | 7 (2.1%) | 14 (4.3%) | 32 (9.8%) | 29 (8.9%) | 0 |
| Other | Rural (N=36) | 0 | 0 | 0 | 0 | 4 (11.1%) | 0 | 0 | 0 |
| | Developing (N=39) | 3 (7.7%) | 1 (2.6%) | 0 | 0 | 4 (10.3%) | 1 (2.6%) | 1 (2.6%) | 0 |
| | Urban (N=3) | 0 | 0 | 0 | 0 | 0 | 0 | 0 | 0 |
| | Total (N=78) | 3 (3.8%) | 1 (1.4%) | 0 | 0 | 8 (10.3%) | 1 (1.3%) | 1 (1.3%) | 0 |

**Supplemental Table 4.** Number and proportion (%) of rodents positive for one or more pathogen PCR assays. A zero indicates no animals were positive, and a – indicates that no animals were tested. The total number of each species tested is given next to each location in parentheses.

| **Species** | | **Orthohantavirus** | **Mammarenavirus** | **Orthohepevirus** | **Paramyxovirus** | ***Betacoronavirus*** | ***T. gondii*** | ***Bartonella* spp.** | ***Leptospira* spp.** |
|---|---|---|---|---|---|---|---|---|---|
| *R. rattus* R3 | Rural (N=30) | 0 | 0 | 0 | 0 | 4 (13.3%) | 1 (3.3%) | 12 (40%) | 3 (10.0%) |
| | Developing (N=61) | 0 | 0 | 7 (11.5%) | 1 (1.6%) | 0 | 7 (11.5%) | 22 (36.1%) | 29 (47.5%) |
| | Urban (N=74) | 3 (4.1%) | 1 (1.4%) | 5 (6.8%) | 2 (2.7%) | 1 (1.4%) | 10 (13.5%) | 45 (60.8%) | 38 (51.4%) |
| *R. rattus* tanezumi | Rural (N=0) | - | - | - | - | - | - | - | - |
| | Developing (N=6) | 0 | 0 | 0 | 0 | 0 | 1 (16.7%) | 2 (33.3%) | 2 (33.3%) |
| | Urban (N=5) | 0 | 0 | 0 | 0 | 0 | 1 (20.0%) | 2 (40.0%) | 1 (20.0%) |
| *R. tiomanicus* | Rural (N=0) | - | - | - | - | - | - | - | - |
| | Developing (N=8) | 0 | 0 | 1 (12.5%) | 0 | 0 | 0 | 5 (62.5%) | 2 (25.0%) |
| | Urban (N=0) | - | - | - | - | - | - | - | - |
| *R. exulans* | Rural (N=2) | 0 | 0 | 0 | 0 | 0 | 0 | 0 | 0 |
| | Developing (N=1) | 0 | 0 | 0 | 0 | 0 | 0 | 0 | 0 |
| | Urban (N=0) | - | - | - | - | - | - | - | - |
| *S. muelleri* | Rural (N=22) | 0 | 0 | 0 | 0 | 0 | 0 | 14 (63.6%) | 4 (18.2%) |
| | Developing (N=41) | 0 | 0 | 0 | 1 (2.4%) | 3 (7.3%) | 1 (2.4%) | 41 (100.0%) | 6 (14.6%) |
| | Urban (N=37) | 3 (8.1%) | 0 | 1 (2.7%) | 2 (5.4%) | 0 | 2 (5.4%) | 32 (86.5%) | 13 (35.1%) |
| *M. whiteheadi* | Rural (N=11) | 0 | 0 | 0 | 0 | 0 | 0 | 0 | 1 (9.1%) |
| | Developing (N=1) | 0 | 0 | 0 | 0 | 0 | 0 | 1 (100.0%) | 0 |
| | Urban (N=0) | - | - | - | - | - | - | - | - |
| *M. ochraceiventer* | Rural (N=2) | 0 | 0 | 0 | 0 | 0 | 0 | 0.0 | 1 (50.0%) |
| | Developing (N=0) | - | - | - | - | - | - | - | - |
| | Urban (N=0) | - | - | - | - | - | - | - | - |
| *N. cremoriventer* | Rural (N=10) | 0 | 0 | 0 | 0 | 0 | 0 | 4 (40.0%) | 0 |
| | Developing (N=4) | 0 | 0 | 0 | 0 | 0 | 0 | 1 (25.0%) | 0 |
| | Urban (N=0) | - | - | - | - | - | - | - | - |

### SUPPLEMENTAL MATERIAL – FIGURES

**Supplemental Figure 1. Results of LDAs**

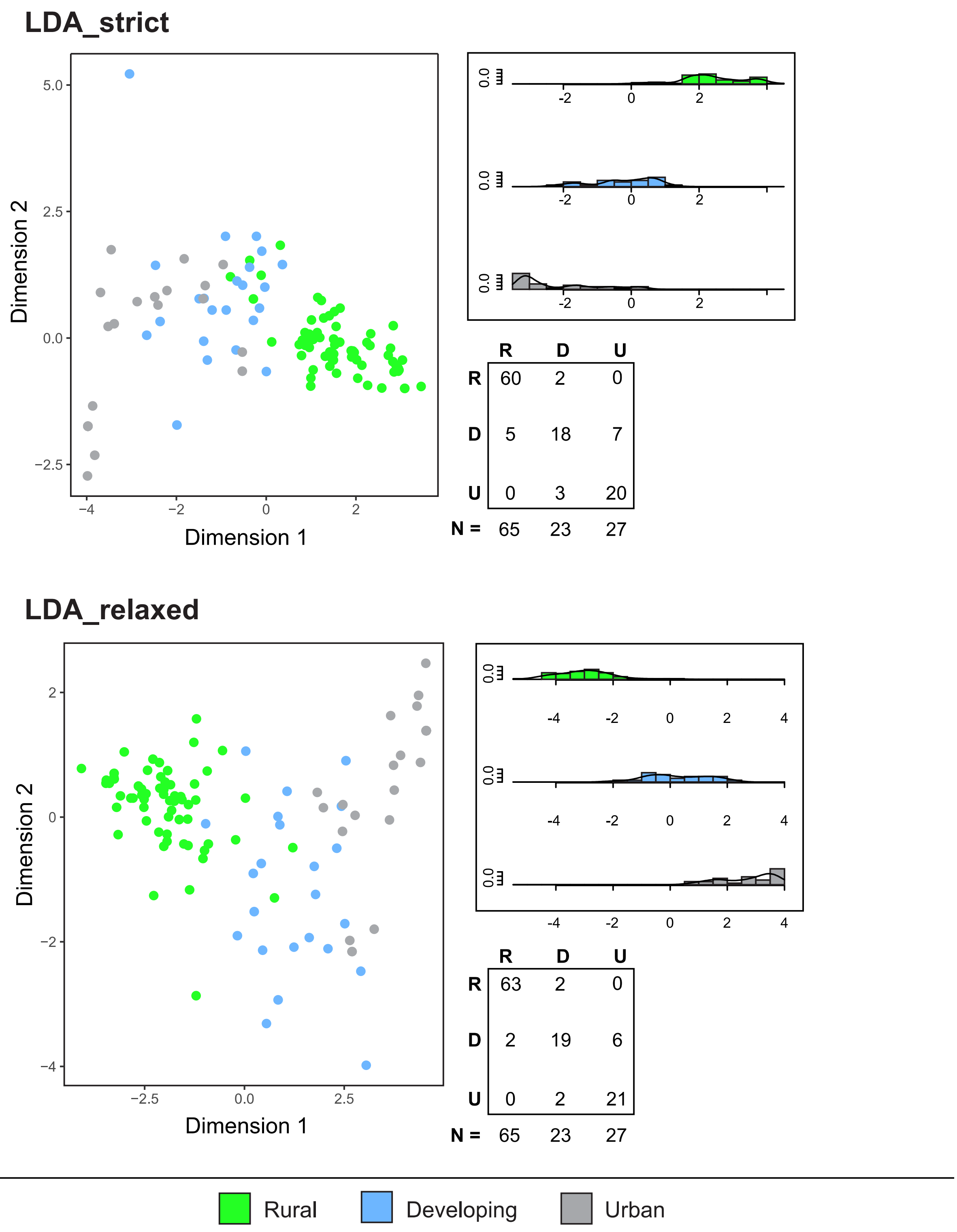

**Supplemental Figure 2. Maximum likelihood phylogeny of the betacoronaviruses identified in this study.**

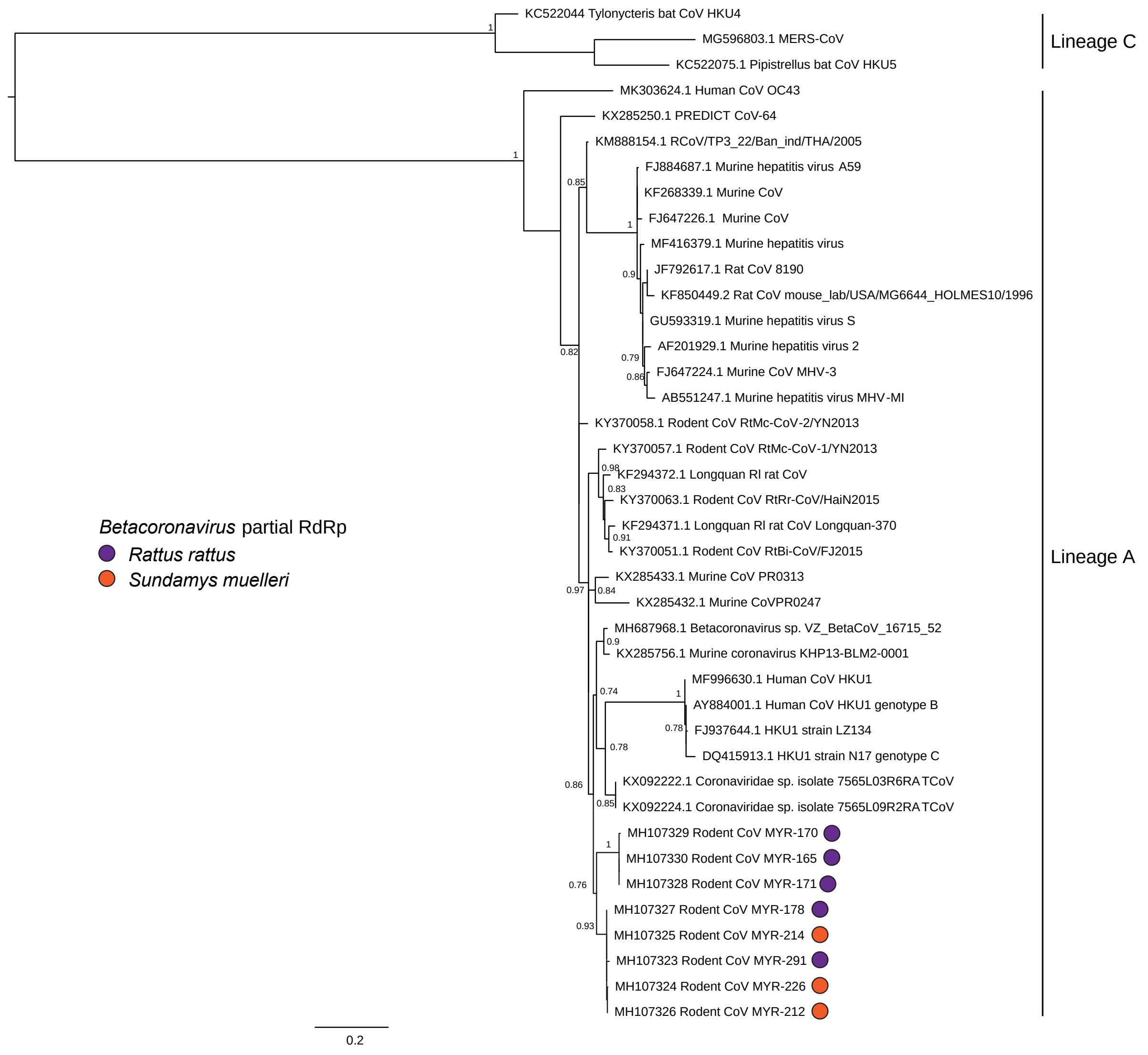

**Supplemental Figure 3. Maximum likelihood phylogeny of the paramyxoviruses identified in this study.**

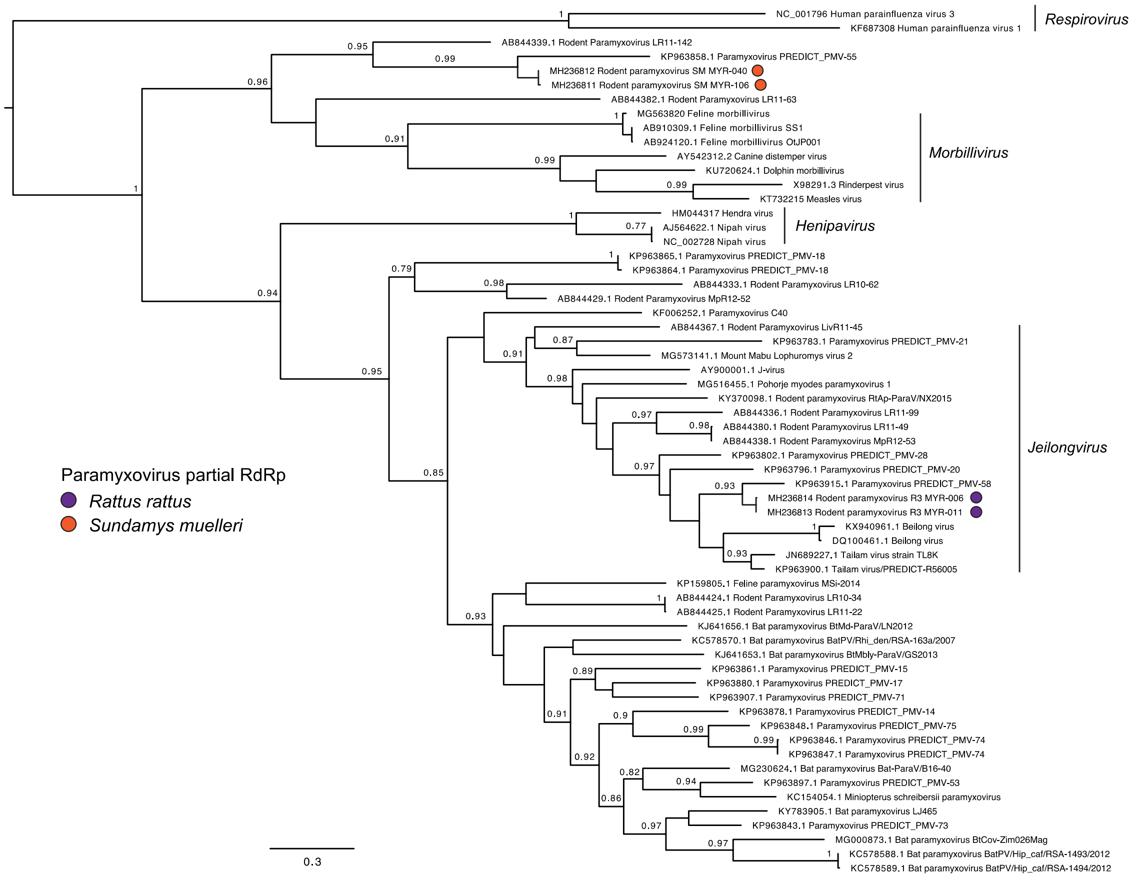
